# A type IVB secretion system contributes to the pathogenicity of *Yersinia pseudotuberculosis* strains responsible for the Far East scarlet-like fever

**DOI:** 10.1101/2024.06.14.598817

**Authors:** Marion Lemarignier, Cyril Savin, Inés Ruedas Torres, Anne Derbise, Charles Coluzzi, Julien Burlaud-Gaillard, Julien Madej, Rémi Beau, Philippe Roingeard, Pierre Lechat, Eduardo Rocha, Jaime Gomez-Laguna, Javier Pizarro-Cerdá

**Affiliations:** Institut Pasteur, Université Paris Cité, CNRS UMR 6047, Yersinia Research Unit, Paris, France; Institut Pasteur, Université Paris Cité, Yersinia National Reference Laboratory, WHO Collaborating Research & Reference Centre for Plague FRA-146, Paris, France; United Kingdom Health Security Agency (UKHSA), Porton Down, Salisbury SP4 0JG, UK; Institut Pasteur, Université de Paris Cité, CNRS UMR 3525, Microbial Evolutionary Genomics, Paris, France; INSERM U1259, Université de Tours et CHRU de Tours, IBiSA Electron Microscopy Platform, Tours, France; Institut Pasteur, Université Paris Cité, Bioinformatics and Biostatistics Hub, Paris, France; Department of Anatomy and Comparative Pathology and Toxicology, University of Córdoba, Spain

**Keywords:** *Yersinia pseudotuberculosis*, Far East scarlet-like fever, pVM82, type IVB secretion system (T4BSS), *dot/icm* genes, virulence factor

## Abstract

*Yersinia pseudotuberculosis* is a food-borne pathogen responsible for a self-limiting gastrointestinal disease in humans known as mesenteric lymphadenitis. A phylogenetically distinct *Y. pseudotuberculosis* cluster from lineages 1 and 8 is associated to a specific syndrome called the Far East scarlet-like fever (FESLF), characterized by skin rash, hyperemic tongue and desquamation. Genome sequencing of FESLF strains previously revealed the presence in the plasmid pVM82 of *dot/icm* genes, homologous to those known to encode a T4BSS in the intracellular pathogens *Legionella pneumophila* and *Coxiella burnetii.* In the present article, we characterized the genomic features and functionality of the *Y. pseudotuberculosis* T4BSS (*y*T4BSS). We found higher *dot/icm* gene identity between *Y. pseudotuberculosis* and *Pseudomonas putida* genes than with those of *L. pneumophila* or *C. burnetii*. We validated the presence of all essential *dot/icm* genes required for the structure of a T4BSS. We then evaluated the conditions required for *y*T4BSS gene expression *in vitro* and identified an influence of temperature, with higher expression at 37°C, which mimicks the mammalian host temperature. The *y*T4BSS is also expressed *in cellulo* during the *Y. pseudotuberculosis* intracellular life cycle and *in vivo* during mouse infection. Although T4BSS functions are well characterized in the intracellular life cycle of *L. pneumophila* and *C. burnetii*, the *y*T4BSS appears to not be required for the intracellular survival nor for the establishment of a replication niche within cells of *Y. pseudotuberculosis*. Interestingly, the *y*T4BSS is implicated in *Y. pseudotuberculosis* FESLF strain pathogenicity when orally inoculated to mice but not during intravenous inoculation. Despite a role in virulence during oral infection, the *y*T4BSS does not influence organ colonization. However, the *y*T4BSS appears to be implicated in induction of important necrosis lesions in mesenteric lymph nodes and cæca of mice. Cytokine profil analyses revealed an induction of production of innate immunity related cytokines and chemokines depending on the *y*T4BSS *in cellulo* using a mouse bone marrow-derived macrophages infection model. Thus, the *y*T4BSS modulates cytokine responses of the host innate immune system during oral infection. In conclusion, the *y*T4BSS is a newly characterized virulence factor implicated in pathogenicity of *Y. pseudotuberculosis* strains from lineage 8 responsible for FESLF.

## Introduction

*Y. pseudotuberculosis* is a Gram-negative pathogen responsible for enteric yersiniosis, a food-borne disease characterized by self-limiting mesenteric adenitis, which can evolve to septicemia in patients displaying underlying disorders such as diabetes, hemochromatosis, or thalassemia (Carniel et al., 2006). Infections by *Y. pseudotuberculosis* are rather rare: in France, the National Reference Laboratory ‘Plague and other yersiniosis’ reports yearly less than 1% of infections due to *Y. pseudotuberculosis,* most yersiniosis being attributed to another enteropathogen, *Y. enterocolitica* (Le Guern et al., 2016; 2022). A similar scenario is observed at the European level (European Center for Disease Prevention and Control, 2024). Nevertheless, *Y. pseudotuberculosis* outbreaks have been reported in several locations around the world including Japan (Inoue et al., 1984), Canada (Nowgesic et al., 1999), Finland (Jalava et al., 2004), and New Zealand (Williamson et al., 2016). The first *Y. pseudotuberculosis* clonal outbreak in France was reported in 2020 (Savin et al., 2022).

*Y. pseudotuberculosis* displays several virulence factors: the chromosomal *inv* gene encodes for invasin, a surface protein that allows bacterial binding to β1 integrin receptors predominantly exposed by Microfold (M) cells in intestinal Peyer’s patches (Clark et al., 1998; Isberg and Falkow, 1985; Marra and Isberg, 1997). This interaction triggers bacterial internalization within host cells and invasion of the intestinal tissue, followed by colonization of draining mesenteric lymph nodes. A type III secretion system (T3SS) encoded in the *Yersinia* virulence plasmid (pYV) is then activated and translocates *Yersinia* outer proteins (Yops) into the cytoplasm of macrophages and immune cells, blocking actin polymerization and inhibiting immune signaling responses dependent on mitogen-activated protein kinases (MAPKs), leading to bacterial extra-cellular proliferation (Cornelis, 2002; Seabaugh and Anderson, 2024). Host iron is sequestered via the siderophore yersiniabactin, encoded in the high pathogenicity island (HPI) which is common to other enterobacterial species such as *Escherichia coli* or *Citrobacter jejuni* (Carniel, 2001; Rakin et al., 2012).

Classical enteric yersiniosis is associated to *Y. pseudotuberculosis* strains present in Western Europe and North America (Fukushima et al., 2001). In Eastern Russia and Japan, a disease named the Far East scarlet-like fever (FESLF) has been associated to a restricted subset of *Y. pseudotuberculosis* strains which have been the cause of two main epidemics reported in 1958 in Vladivostock, Russia and in 1977 in Kanazawa, Japan (Amphlett, 2016; Fukushima et al., 2001; Tseneva et al., 2012). Besides digestive symptoms, FESLF is a longer infection than classical enteric yersiniosis, characterized by fever, skin rash, hyperemic tongue, ending with desquamation (Amphlett, 2016; Mollaret et al., 1990). The presence of a superantigen, the *Y. pseudotuberculosis*- derived mitogen (YPMa), has been reported in Far Eastern *Y. pseudotuberculosis* strains and this virulence factor might be releated to the systemic symptoms detected in FESLF patients (Abe et al., 1997; Abe et al., 2003; Fukushima et al., 2001; Ueshiba et al., 1998). The genome sequencing of a *Y. pseudotuberculosis* FESLF strain identified the presence of a plasmid (pVM82), which had been previously associated to bacterial pathogenicity and to modified cellular immune responses in infected hosts (Dubrovina et al., 1999; Gintsburg et al., 1988; Sever et al., 1991). Interestingly, in the pVM82, Eppinger et al., 2007 identified *dot/icm* (*defect in organelle trafficking*, *intracellular multiplication*) genes known to encode a type IVB secretion system (T4BSS).

T4BSS are typically represented by the effector translocator Dot/Icm system from intracellular pathogens such as *Legionella pneumophila* and *Coxiella burnetii* (Berger and Isberg, 1993; Brand et al., 1994; Segal and Shuman, 1997). However T4BSS also include conjugative system members from the plasmid incompatibility group I such as the R64 plasmid from *Salmonella enterica* serovar Typhimurium and the ColIb-P9 plasmid from *Shigella flexneri* (Hartskeerl et al., 1985; Sampei et al., 2010) and recently a bacterial killing system from *Pseudomonas putida* (Purtschert-Montenegro et al., 2022). The T4BSS of *L. pneumophila* and *C. burnetii* is an essential virulence factor for their intracellular life cycle, secreting a wide range of effectors involved in the manipulation and disruption of cellular organelles, host metabolism, cell death and immune responses (Lührmann et al., 2017; Qiu and Luo, 2017). The Dot/Icm T4BSS system from *L. pneumophila* has the most characterized structure to date and consists of a transport channel that spans the bacterial envelope for substrates translocation, composed of an outer membrane complex core (OMCC: DotC, DotD, DotH), a central cylinder channel connecting the outer and inner membranes (DotG, DotF, IcmX, IcmF, DotA), inner membrane proteins (DotJ, IcmT, IcmV, DotI, DotU, DotE) and a cytoplasmic complex of ATPases (DotO, DotB) (Chetrit et al., 2018; Ghosal et al., 2019; Macé et al., 2022b). A type IV coupling complex (T4CC) acts as a substrate delivery platform for the transport channel which is composed of an ATPase (DotL), adaptor proteins (DotM, DotN, DotY, DotZ) with chaperones (IcmS, IcmW, LvgA) (Macé et al., 2022a).

Here, we studied the T4BSS in a *Y. pseudotuberculosis* FESLF strain (*y*T4BSS). We validated *y*T4BSS gene expression *in vitro*, *in cellulo* and *in vivo*, observing a higher expression rate at 37°C, mimicking a mammalian host environment. We studied the function of the *y*T4BSS and did not detect its implication in the intracellular life cycle of *Y. pseudotuberculosis*, but we found it to be a virulence factor involved in bacterial pathogenicity in orally infected mice. The *y*T4BSS appears to modulate the cytokine innate immune responses and is associated to necrotic lesions in infected organs.

## Results

### Distribution, organization and genomic features of the *y*T4BSS gene cluster

The genome sequencing of the FESLF-causing *Y. pseudotuberculosis* reference strain IP31758 (Eppinger et al., 2007) identified the presence of a *y*T4BSS in the pVM82 plasmid. The Eppinger *et al*. study also revealed in the strain IP31758, the absence (probably lost in the laboratory environment) of the pYV plasmid encoding a T3SS, fully required for virulence in pathogenic *Yersinia* spp. (Cornelis, 2002). Since the absence of pYV would have precluded any virulence studies in cellular or animal infection models, we decided to use a *Y. pseudotuberculosis* isolate closely related to the IP31758 strain in order to investigate the functionalities of the *y*T4BSS. We chose therefore to use in our study the strain SP-1303, which: a) is genetically very close to the strain IP31758 and belongs to the *Y. pseudotuberculosis* lineage 8, which harbors strains associated to FESLF (Savin et al., 2019), and b) it has both the pYV and the pVM82 (Lemarignier et al., 2023) (Figure 1A).

**Figure 1:**
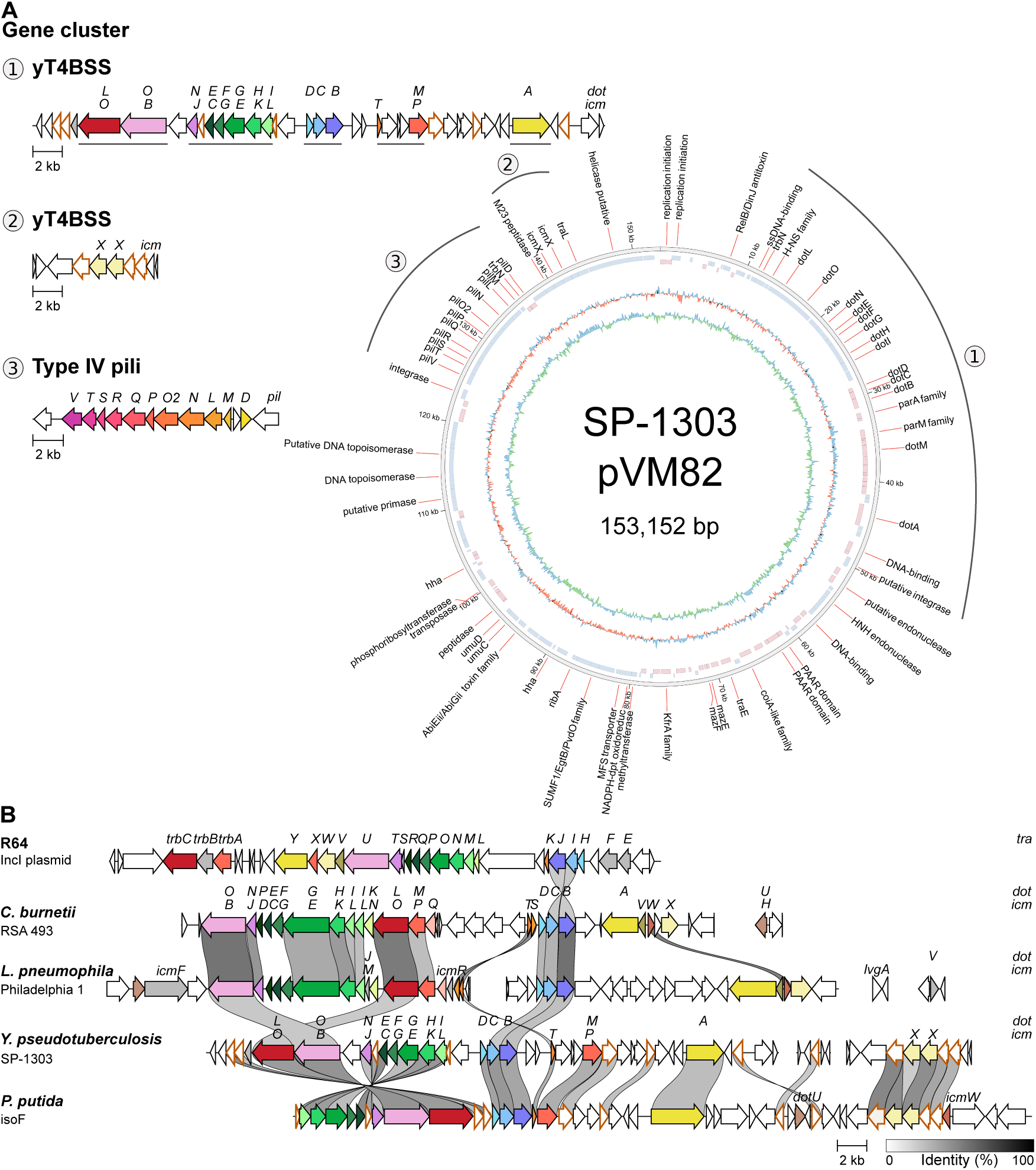
Gene organization and homology of the *y*T4BSS gene cluster in pVM82 plasmid from *Y. pseudotuberculosis* strain SP-1303. **(A)** Plasmid map of the pVM82 from *Y. pseudotuberculosis* FESLF strain SP-1303 with annotated genes product. The first outer ring corresponds to gene products, without labeling hypothetical protein products. The second ring represents gene position (blue: positive or red: negative strand). Third ring shows GC skew. Inner ring shows GC content variation from the mean. Figure generated with Circos from the annotated Genbank sequence of pVM82 (ID: CP130902.1) (Krzywinski et al., 2009). The structure of the *y*T4BSS and of a type IV pili gene clusters are further highlighted. **(B)** T4BSS gene clusters conservation and comparison between *Y. pseudotuberculosis* SP-1303 (GenBank accession no. NZ_CP130902.1), the IncI plasmid R64 (NC_005014), *Coxiella burnetii* RSA 493 (NC_002971), *Legionella pneumophila* Philadelphia 1 (NC_002942), *Pseudomonas putida* isoF (CP072013.1) with a minimum alignement sequence identity of 30 %. Genes coloured white are not T4BSS genes and those with brown outlines share sequence identity. Genes coloured in light gray are strain specific. Figure generated with clinker (Gilchrist and Chooi, 2021).

The *y*T4BSS gene cluster of the strain SP-1303 is genetically similar to the one previously described in the strain IP31758: it displays several *dot/icm* genes which are homologous to the T4BSS genes present in the bacterial intracellular pathogens *L. pneumophila, C. burnetii* and the recently T4BSS described in the soil saprophyte *P. putida* (strain isoF) (Figure 1B) (Purtschert-Montenegro et al., 2022). 13 *dot/icm* genes (out of 27 in *L. pneumophila* and 23 in *C. burnetii*) were first identified by Eppinger et al., 2007 and are known to encode essential structural components of the translocon apparatus, including the crucial cytoplasmic ATPases (*dotO/icmB* and *dotB*) as well as encoding key elements of the T4CC. By using Macsyfinder (Abby et al., 2014), we were able to identify three additional *dot/icm* genes on the pVM82, also present in *L. pneumophila*, *C. burnetii*, and *P. putida*: *icmT,* encoding a structural cytoplasmic membrane protein (Bitar et al., 2005), as well as two copies of *icmX (* annotated: *icmX1* and *icmX2)*, known to encode a plug structural periplasmic protein (Matthews and Roy, 2000) which were not initially recognized as such by Eppinger et al., 2007 (Figure 1A). The two copies of *icmX* are encoded in a different operon, in close proximity to a type IV pili gene cluster on the pVM82 (Figure 1A). Some *dot/icm* genes shared by *L. pneumophila* and *C. burnetii,* implicated in the structure and function of their T4BSS are absent in the *y*T4BSS such as genes encoding the chaperones *icmS* and *icmW* (Cambronne and Roy, 2007; Macé et al., 2022a; Xu et al., 2017), genes encoding the inner membrane structural proteins *icmV*, *icmQ*, *dotU/icmH*, *dotP/icmD,* as well as the genes *dotK/icmN* forming the α-density of the OMCC (Durie et al., 2020; Ghosal et al., 2019). The genes *dotZ*, *dotY, lvgA, icmR, dotJ/icmM, icmF* and *dotV* were also not found in the *y*T4BSS gene cluster but are known to be specific to *L. pneumophila* (Zamboni et al., 2003; Zusman et al., 2003). Interestingly, higher percentage of sequence identity with a greater number of homologous *dot/icm* genes were found between the *y*T4BSS gene cluster and the one from *P. putida* strain isoF compared to those of *L. pneumophila* and *C. burnetii* (Figure 1B).

Several T4BSS have been described as required for conjugation, implicating the function of an essential relaxase which binds to the origin of transfer (*oriT*) sequence, resulting in ssDNA transfer to the T4BSS (Cruz et al., 2010). By running Macsyfinder (Abby et al., 2014) and oriTfinder (Li et al., 2018) on the pVM82 of *Y. pseudotuberculosis* SP-1303, no *oriT* and no relaxase were identified, suggesting that the *y*T4BSS is not implicated in conjugation of the pVM82. In addition, very low identity with few genes is detected between *tra* genes of the representative conjugative T4BSS on the plasmid R64 and the *y*T4BSS gene cluster (Figure 1B).

To have a broader view of the presence of the *y*T4BSS in all lineages from the species *Y. pseudotuberculosis*, we investigated the distribution of *dot/icm* genes in 287 bacterial genomes (Savin et al., 2019). The full *y*T4BSS gene cluster was only observed in the majority of strains from the ancestral lineage 8 (Figure S1A). Incomplete *y*T4BSS were observed in strains from lineages 6 and 11 (Figure S1A). Interestingly, the clinical isolates of *Y. pseudotuberculosis* responsible for FESLF are grouped in lineages 1 and 8, suggesting that the *y*T4BSS could participate in the pathogenicity of FESLF strains. Of note, the genes encoding the superantigen YPMa are also present in lineages 1 and 8, but it is also present in lineages 23 and 29 which have not been associated with FESLF. In order to investigate whether the *y*T4BSS gene organization is conserved in *Y. pseudotuberculosis* strains from lineage 8, we analyzed the genomes of 46 isolates from this bacterial group. As shown in Figure S1B, the gene organization of the *y*T4BSS gene cluster is highly similar between strains with a conserved synteny. The presence of the main essential structural and functional *dot/icm* genes (discussed above) and the conserved synteny in multiple strains suggest that the *y*T4BSS is not in degradation mode and could be functional (Nagai and Kubori, 2011).

Finally, we explored in the NCBI database using cblaster (Gilchrist and Chooi, 2021) to investigate whether the *y*T4BSS is also found in other genomes of the genus *Yersinia*. We were able to identify complete *dot/icm* gene clusters with high homology (>80% identity and >95% coverage) only in pathogenic species, including strains from the human enteropathogen *Y. enterocolitica* as well as strains from the fish pathogen *Y. ruckeri* but not in *Y. pestis* (Figure S2). Incomplete isolated clusters of genes were found in the environmental species *Y. alsatica* as well as in other strains of *Y. enterocolitica* and *Y. ruckeri.* The presence of closely homologous *y*T4BSS in phylogenetically distant strains of *Yersinia* spp. (*Y. enterocolitica*, *Y. ruckeri)* suggests an independent acquisition.

In conclusion, *Y. pseudotuberculosis* strains from lineage 8, associated with FESLF, harbor a complete *y*T4BSS gene cluster encoded on the pVM82 plasmid, highly similar to known T4BSS in *L. pneumophila, C. burnetii* and *P. putida,* and it might be implicated in the development of FESLF.

### The *y*T4BSS is expressed *in vitro*, *in cellulo* and *in vivo*

To address the functionality of the *y*T4BSS, we first investigated its putative expression under different conditions *in vitro, in cellulo* and *in vivo*. We identified in the main *y*T4BSS gene cluster five putative transcriptional units out of the eleven previously characterized in *L. pneumophila* (Sahr et al., 2012) (underlined in Figure 1A). By qRT-PCR, we measured the expression level of one gene of each transcriptional unit (*dotA, dotM, dotB*, *dotG* and *dotO*) in LB media, at post-exponential phase (7-8 h of growth) and at different temperatures (21°C, 28°C and 37°C). For the five analyzed genes, we observed an increase in their expression level between 21°C and 28°C (environmental temperatures with 28°C as optimal growth temperature) but the differences were not statistically significant; importantly, for all the genes we observed a significant increase in expression levels between 21°C and 37°C (host-associated temperature) (Figure 2A). In the case of four genes out of the five (*dotA*, *dotM*, *dotB* and *dotO*), significant differences were also observed between 28°C and 37°C, suggesting overall an increased expression of the *y*T4BSS in conditions mimicking the host environment (37°C).

**Figure 2:**
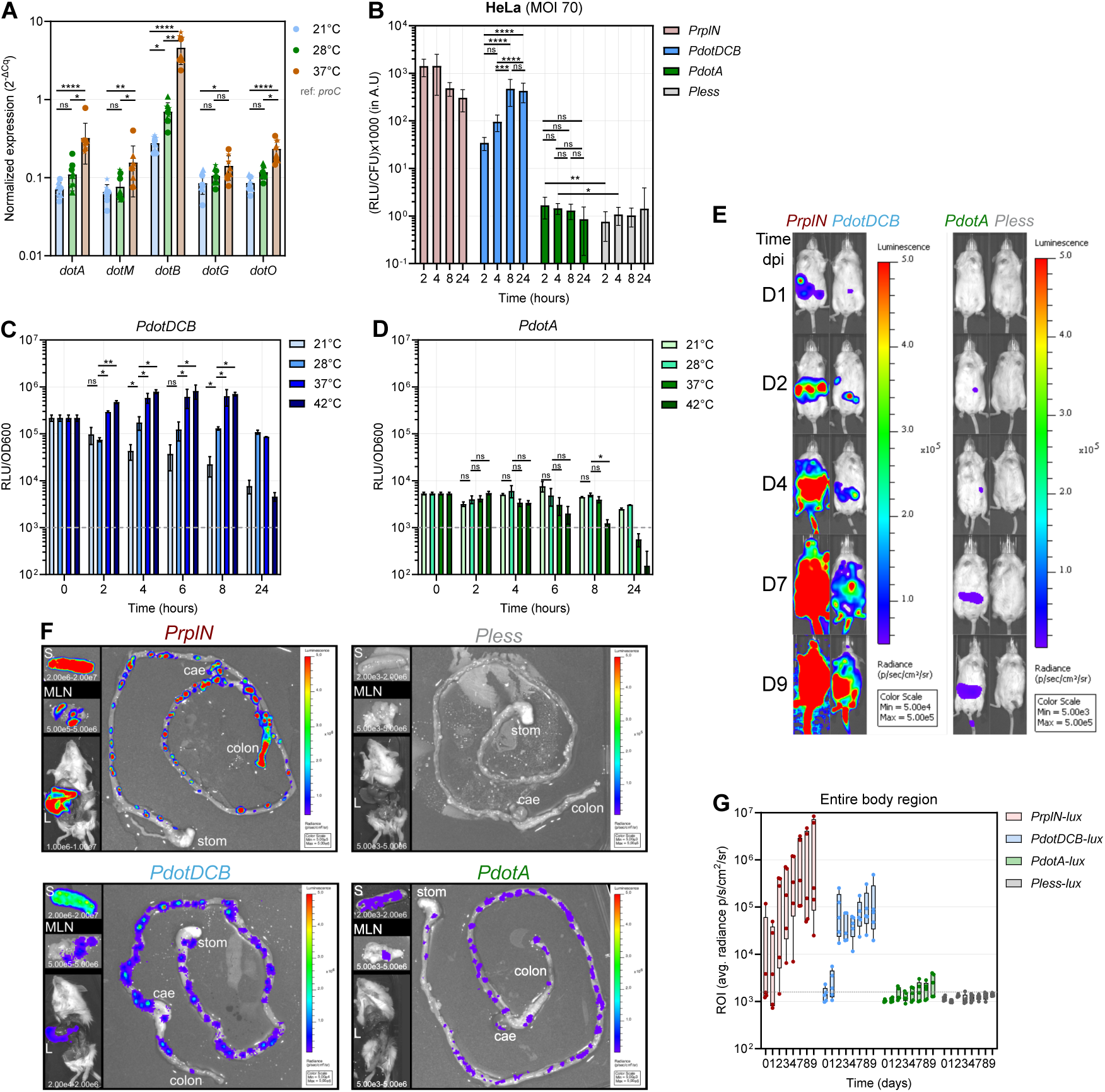
*y*T4BSS genes are expressed *in vitro*, *in cellulo* and *in vivo*. **(A)** Normalized expression of five *y*T4BSS genes to the reference gene *proC*, in the SP-1303 WT strain cultured in LB media at late-exponential phase (7-8 h of growth) under different temperatures (21°C, 28°C, 37°C). Ct obtained by qRT-PCR are normalized to the Ct measured with the housekeeping gene *proC* and plotted as 2^-ΔCq^. Data are representative of 3 independent experiments with means and SD. Kruskal-Wallis tests corrected for multiple comparisons using Dunn’s test were performed on each gene. **(B)** Photon emission measured during the intracellular cycle of bioluminescent reporter strains within HeLa cells. *Y. pseudotuberculosis* SP-1303 strains harboring different bioluminescent reporters were grown O/N at 28°C prior infection for 1 h (MOI of 70) at 37°C, followed by gentamicin treatment (+1 h, +3 h, +7 h, +23 h). RLU measured for each well were normalized by CFUs counted per well. Data shown are representative of three independent experiments with 4 replicates/condition with means and SD. One-way ANOVA test corrected for multiple comparisons using Tukey’s test were performed for each reporter. Unpaired t-tests were performed between *PdotA-lux* and *Pless-lux*. **(C-D)** Measurement of photons emission produced during bacterial growth in LB media, at different temperatures (21°C, 28°C, 37°C, 42°C) and over time for the bioluminescent reporters *PdotDCB-lux* and *PdotA-lux*. Bacteria harboring the reporters were grown O/N at 28°C prior to the experiment. Relative light units (RLU) measured using a Glomax plate reader, were normalized by OD_600nm_. Horizontal gray lines show the background threshold measured with the *Pless-lux* reporter, a construction without promoter inserted. Data are representative of two independent experiments with means and SD. One-way ANOVA tests corrected for multiple comparisons using Tukey’s tests were performed at each time point. **(E)** One representative example of OF1 mice bread-feeded and inoculated with the *Y. pseudotuberculosis* SP-1303 strains harboring bioluminescent reporters (inoculum: 6*×*10^8^ CFU/mouse), monitored each day until day 9 post-infection (n=5/condition) using an IVIS Spectrum system. The same radiance scale (in p/sec/cm^2^/sr) was used for the high signal found in infected conditions with *PrplN-lux* and *PdotDCB-lux* strains, whereas another radiance scale is used for *PdotA-lux* and *Pless-lux*. Photon emission is plotted in graph D. **(F)** Dissection of mice infected with *Y. pseudotuberculosis* SP-1303 strains harboring bioluminescent reporters at day 9 post-infection. Stom: stomach, cae: cæcum, S: spleen, MLNs: mesenteric lymph nodes, L: liver. Radiance scales used in p/sec/cm^2^/sr are indicated for each organ. **(G)** Photon emission measurements over time, in the region of interest (ROI) of mice orally infected (inoculum: 6*×*10^8^ CFU/mouse) with *Y. pseudotuberculosis* SP-1303 strains harboring different bioluminescent reporters (*PrplN-lux*, *PdotDCB-lux*, *PdotAlux*, *Pless-lux*). ROIs covering the entire mouse body were drawn using the Living Image v.4.5 software. Data represent a single experiment with 5 mice/condition.

We then complemented these *in vitro* results by investigating the activity of the promoter regions of the highly conserved T4BSS genes *dotDCB* and *dotA* upon fusion to the *luxCDABE* operon of *Photorabdus luminescens* (Derbise et al., 2019), monitoring bioluminescence production in conditions similar as those tested above by qRT-PCR. Both promoters *PdotDCB* and *PdotA* were activated at the four tested temperatures 21°C and 28°C (environmental conditions), 37°C and 42°C (mimicking host-associated basal and high fever temperatures, respectively), when compared with the background signal of a construction without promoter (*Pless*), all along bacterial growth (Figure 2C-D, Figure S3). Interestingly, in contrast to *PdotA* with a basal constant activity close to background noise, regardless of the temperature, *PdotDCB* activation is temperature dependent: from an overnight (O/N) preculture at 28°C, a one log10 decrease in normalized photon emission was measured when bacteria were newly grown at 21°C during 24 h; on the contrary, when newly grown at 37°C or 42°C, a significant increase in *PdotDCB* activity was measured over time (Figure 2C). Altogether, these results suggest that the *y*T4BSS is expressed *in vitro* during bacterial growth and in a temperature-dependent manner, with higher expression at temperatures mimicking the mammal host (37°C) than in environmental ones (21°C, 28°C).

Because the T4BSS of *L. pneumophila* and *C. burnetii* is essential for their intracellular life cycle (Qiu and Luo, 2017), the activity of the *PdotDCB* and *PdotA* bioluminescent reporters was evaluated upon infection of epithelial human HeLa cells. After 1 h of infection, gentamicin was added, left for bacterial extracellular killing for additional time periods (+1 h, +3 h, +7 h and +23 h) and photon emissions were monitored and normalized by CFUs per well. *PdotA* was active *in cellulo* at 2 h and 4 h post-infection (pi) when compared to the background noise of the *Pless* construction, but the signal was not significantly different from the background at 8 hpi and 24 hpi (Figure 2B). To the contrary, *PdotDCB*, also pregrown at 28°C, is upregulated all along the course of intracellular life cycle within HeLa cells, suggesting that temperature and/or cellular environments are major drivers of *y*T4BSS expression.

We finally investigated *in vivo* the spatio-dynamic expression of the *y*T4BSS using a murine oral infection model (Derbise et al., 2020). Beside the *Y. pseudotuberculosis* SP-1303 strains harboring the *PdotA* and *PdotDCB* inductible reporters as well as the control *Pless* construct described above, mice were also infected with a strain harboring a constitutive promoter controlling bioluminescence production (*PrplN*) (Figure 2E-G). The *PdotDCB* reporter was found to be active all along the course of mice infection, characterized first by bioluminescence signals in the abdominal region and later reaching the mouse tail, the latter corresponds to a septicemic stage with bacteria present in the blood stream (Figure 2E). The *PdotA* reporter displayed a similar pattern as *PdotDCB* but its signal was very low, close to the background noise defined with *Pless*-infected mice (Figure 2E). It is interesting to notice that the quantification of the bioluminescence signal in the entire body of each mouse reveals a high increase in photons emission for *PdotDCB*-infected mice between day 1 and 2 pi, as compared with a linear signal increase in *PrplN* -infected mice correlated with bacterial replication, suggesting an up-regulation of *PdotDCB* at early times pi (Figure 2G). In addition, the *PdotDCB* and *PdotA* are active in all infected organs: all along the gastrointestinal tract (in the intestinal lumen, Peyer’s patches, cæcum and colon) as well as in the spleen, pool of mesenteric lymph nodes (MLNs) and liver (exception for *PdotA*) (Figure 2F). Thus, *y*T4BSS is active *in vivo*, all along the course of mouse infection without any organ specificity.

In conclusion, the *y*T4BSS is active and expressed *in vitro* in a temperature-dependent manner, with increased expression *in vitro* at 37°C (a temperature mimicking mammalian host body), during the intracellular life cycle of *Y. pseudotuberculosis* in mammalian cell lines, as well as during mouse infection in all infected organs.

### The *y*T4BSS is not implicated in the intracellular life cycle of *Y. pseudotuberculosis*

*Y. pseudotuberculosis* is able to invade host cells via interactions between its surface molecule invasin and host cell β1-integrin receptors (Isberg et al., 1987). Within epithelial cells but also macrophages, *Y. pseudotuberculosis* has been reported to proliferate in autophagosomes (Ligeon et al., 2014; Moreau et al., 2010) but the bacterial virulence factors implicated in the biogenesis of this intracellular replication niche have not been identified so far (Lemarignier and Pizarro-Cerdá, 2020). Since *L. pneumophila* and *C. burnetii* use their respective T4BSS to generate intracellular replication vacuoles via hijacking several host functions including membrane trafficking (Christie et al., 2017; Lockwood et al., 2022; Qiu and Luo, 2017), we investigated whether the *y*T4BSS was implicated in modulating the *Y. pseudotuberculosis* intracellular behavior. For that, we generated in the strain SP-1303 a single deletion mutant of the gene *dotB*, encoding a crucial cytoplasmic ATPase, since *dotB* has been shown to be essential for the functionality of the T4BSS in *L. pneumophila* and *C. burnetii* (Sexton et al., 2004; 2005; Zamboni et al., 2003) and we showed in the precedent section that the *dotB* gene is upregulated *in cellulo* and *in vivo* upon bacterial infection.

Using gentamicin invasion assays, we first investigated in HeLa cells the intracellular survival of WT versus Δ*dotB* mutant bacteria upon different infection times (Figure 3A). As shown in Figure 3A, no significant differences in intracellular bacterial loads were observed between the WT strain and the Δ*dotB* mutant at all investigated time points (2, 4, 8, 24 hpi). Similar results were found when studying infection of murine RAW264.7 macrophages, of mouse bone marrow-derived macrophages (mBMDM) or human blood monocyte-derived macrophages (Figure S4, Figure S5), suggesting that the *y*T4BSS is not implicated in cellular invasion nor modulation of the intracellular survival of *Y. pseudotuberculosis*.

**Figure 3:**
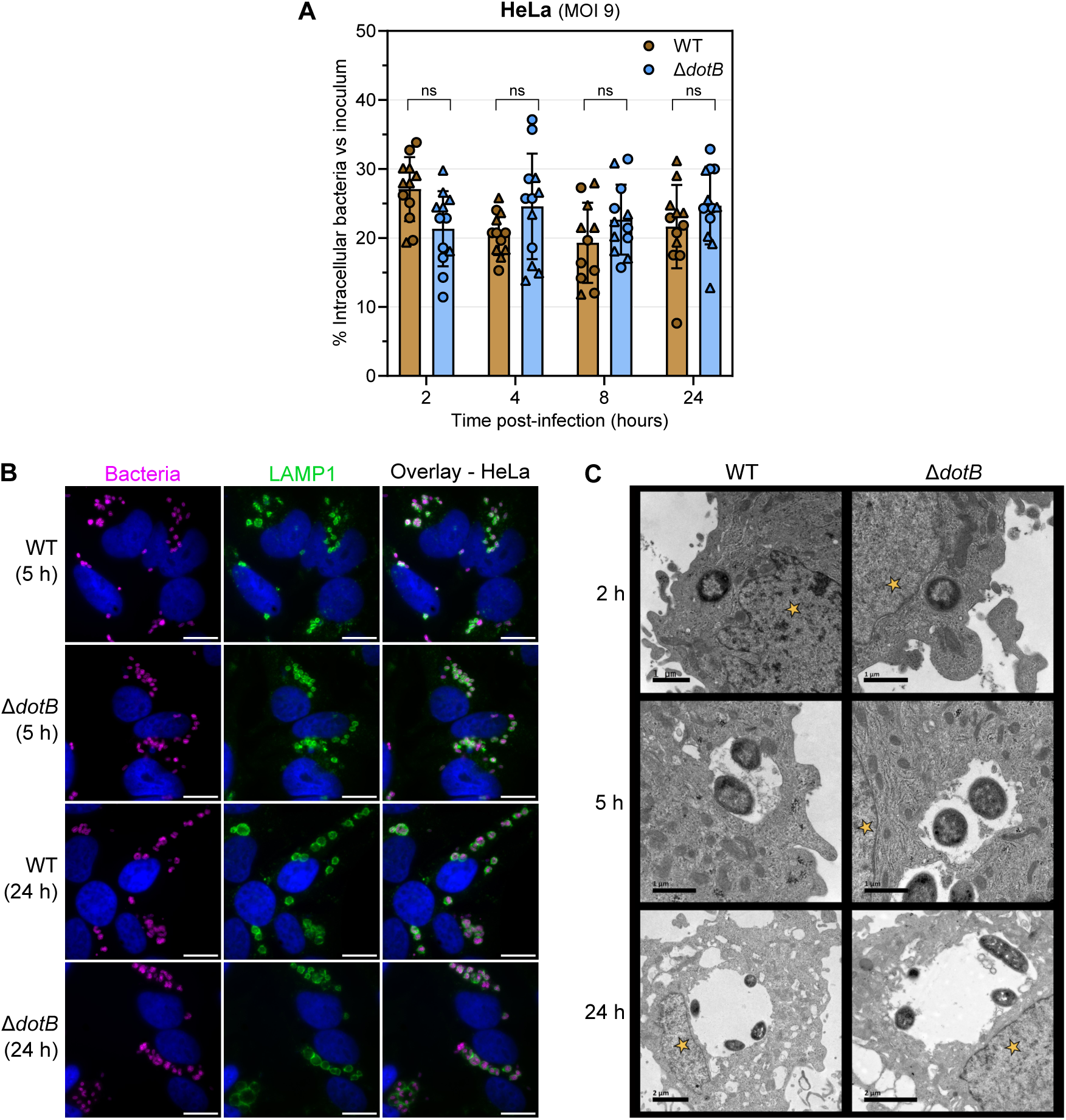
The *y*T4BSS does not contribute to the intracellular life cycle of *Y. pseudotuberculosis* within epithelial HeLa cells. **(A)** Intracellular survival of WT and Δ*dotB* mutant strains within HeLa cells (MOI 9) at different time post-infection (2 hpi, 4 hpi, 8 hpi, 24 hpi). Data are representative of two independent experiments (triangle and round differentiate the experiments) with means and SD. Multiple unpaired t tests were performed. **(B)** Representative images of HeLa cells infected (MOI 25) with the WT or Δ*dotB* strains at 5 hpi or 24 hpi by fluorescent microscopy using antibodies against the surface O-antigen (O1b) of *Y. pseudotuberculosis* (Cy5, magenta), antibodies against LAMP1 (FITC, green) and Hoechst (blue). Scale bars are 10 µm. Overlay images were generated using the ImageJ software. **(C)** Representative images of vacuolar structures within HeLa infected cells with the WT or Δ*dotB* strains at different time (2 hpi, 5 hpi, 24 hpi) using transmission electron microscopy (TEM). Yellow stars highlight the position of nuclei. Scale bars are 1 µm (2 h, 5 h) and 2 µm (24 h).

We then analyzed by fluorescence microscopy the intracellular behavior of the SP-1303 WT strain and of the Δ*dotB* mutant by infecting mammalian host cells for different periods of time (5 h and 24 h), and labeling bacterial intracellular compartments with antibodies against the lysosomal membrane-associated protein 1 (LAMP-1). In HeLa cells, we were able to observe the formation of *Yersinia-*containing vacuoles (YCVs) labeled with LAMP-1 at 5 hpi, and we noted a significant enlargement of these compartments at 24 hpi (Figure 3B); however, no differences were noted between YCVs generated by the WT versus those formed by the Δ*dotB* mutant. We were also not able to detect differences in their intracellular behavior in epithelial Caco-2 cells, RAW264.7 macrophages, mBMDM or human blood monocyte-DM (Figure S4, Figure S5). To further gain insight into the ultrastructure of the YCVs, we infected HeLa cells for different time points (2, 5 and 24 hpi) and we analyzed them using transmission electron microscopy (TEM). As we previously observed by fluorescence microscopy, in both infected cases (WT or Δ*dotB* mutant), YCVs were clearly visible at 5 hpi, and the size of the vacuoles increased at 24 hpi, reaching around 7 µm (Figure 3C). YCVs were formed by a single membrane vacuole as previously described in HeLa cells (Ligeon et al., 2014) and preferentially localize close to the nucleus. Again, no apparent differences in the ultrastructure were observed between YCVs formed by WT versus Δ*dotB* mutant bacteria, which are both preferentially located in contact with the vacuolar membrane. Thus, the *y*T4BSS does not take part in vacuole biogenesis, maturation and localization within cells.

Our results therefore indicate that, to the opposite of *L. pneumophila* and *C. burnetii,* the *y*T4BSS is not required to modulate the intracellular trafficking of *Y. pseudotuberculosis*.

### The *y*T4BSS is required for *Y. pseudotuberculosis* full virulence in a mouse oral infection model

To investigate the potential contribution of the *y*T4BSS in *Y. pseudotuberculosis* pathogenicity *in vivo*, we infected orally or intravenously BALB/c mice with WT or Δ*dotB* mutant strains and monitored first animal survival (Figure 4A-B). Interestingly, a higher percentage of mice survived upon oral infection with the Δ*dotB* mutant compared to animals challenged with the WT strain, demonstrating an attenuation of the virulence of Δ*dotB* mutant *in vivo* (Figure 4A). A complemented Δ*dotB*::*dotB* strain restored the virulence phenotype in orally infected mice. In contrast, no differences in mice survival was observed with the WT strain or the Δ*dotB* mutant when inoculated intravenously (Figure 4B, Figure 7). Thus, these results demonstrate a contribution of the *y*T4BSS in *Y. pseudotuberculosis* virulence upon oral infection in mice, but not upon intravenous infection.

**Figure 4:**
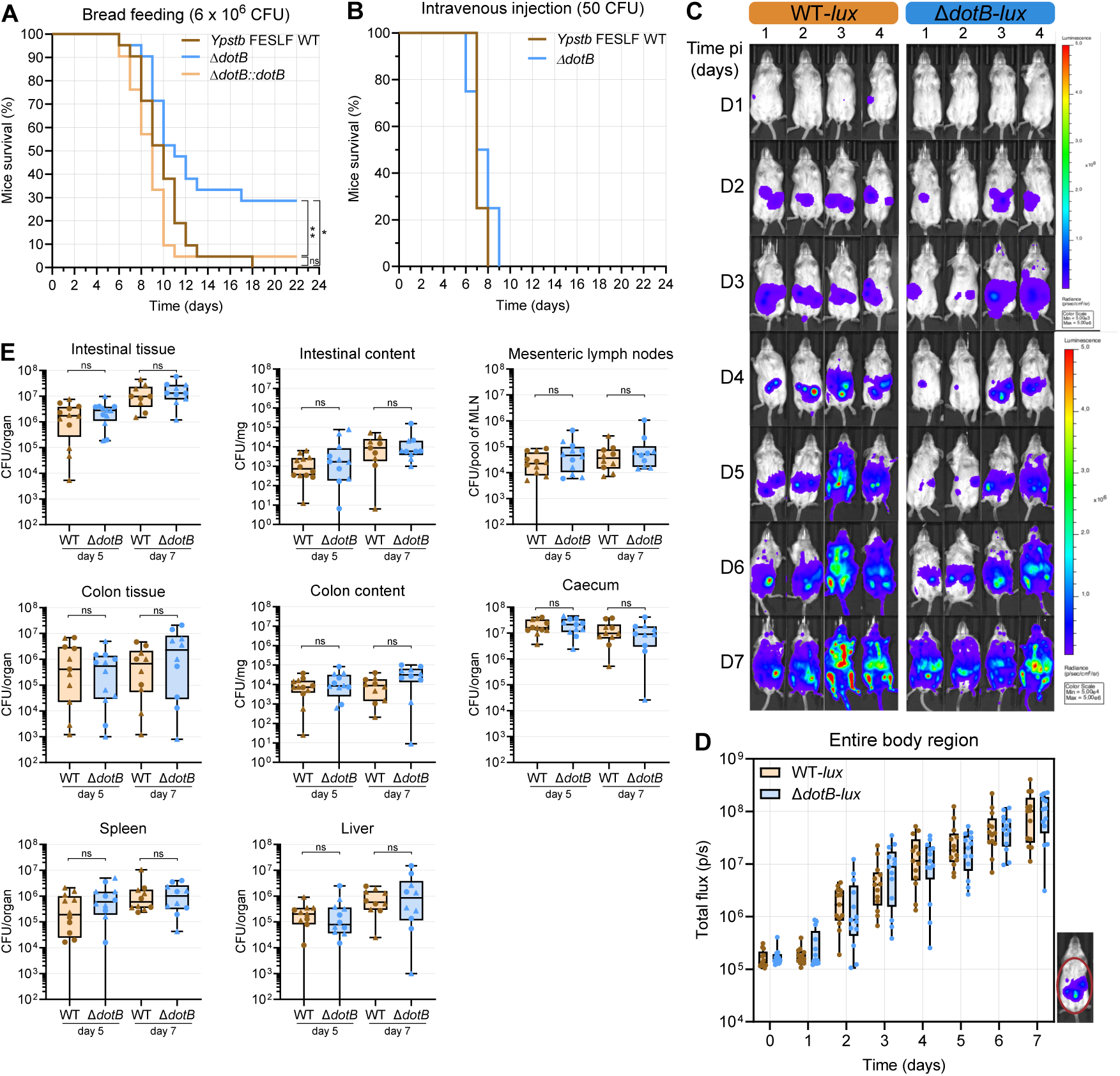
The *y*T4BSS contributes to *Y. pseudotuberculosis* virulence in orally-infected mice without affecting organ colonization. **(A)** Survival monitoring of bread-feeded mice inoculated with WT, Δ*dotB* mutant or Δ*dotB::dotB* complemented strains at 6*×*10^6^ CFU/mouse. Three independent experiments were performed (n=7 mice/condition/experiment). Log- rank (Mantel-Cox) test was performed (WT/Δ*dotB*: p-value=0,0107; Δ*dotB::dotB*/Δ*dotB*: p-value=0,0014). **(B)** Survival monitoring of mice after intravenous injection with 50 CFU/mouse of WT, Δ*dotB* mutant strains. Representation of a single experiment (n=5/condition). **(C)** Photon emission monitoring in mice infected with the constitutive bioluminescent WT*-lux* or Δ*dotB-lux* strains overtime using an IVIS imaging system. Mice number 1, 2 or mice number 3, 4 are representative of respectively slow and rapid kinetic of bacterial colonization. Same radiance scale bar (in p/sec/cm^2^/sr) was used for days 1-3 pi and another for 4-7 pi. **(D)** Total photon flux (p/s) measured on drawn ROIs covering the mouse body using Living Image 4.5 software. Pool of two independent experiment with groups of 6-7 mice. **(E)** Bacterial load per infected organ from orally infected mice (6*×*10^6^ CFU/ mouse) with WT or Δ*dotB* mutant strains. Intestine, colon, cæcum, spleen liver and mesenteric lymph nodes were collected at day 5 and 7 pi and homogenized for CFUs counting. Intestines and colons were longitudinaly oppened for content collection for CFUs, treated for 1 h with gentamicin before homogenization for CFUs counting (intestinal and colon tissues). Data shown are representative of two independent experiments with 5-7 mouse/condition. No statistical differences in bacterial loads were identified

To further gain insight into the mechanisms underlying the reduced virulence of the Δ*dotB* mutant in orally infected mice, we constructed constitutive bioluminescent WT or Δ*dotB* strains and quantified photon emission in orally infected mice each day during infection, in order to obtain information on the kinetics of infection and bacterial localization throughout infection (Figure 4C-D). In both WT and Δ*dotB*-infected groups, starting for few mice at day 1 pi, photons were recorded in the abdominal region with increasing signal intensity over time, suggesting bacterial colonization of the gastrointestinal tract. For some mice in both infected groups, at day 5 or 6 pi signals were detected in the tail, an indication of blood colonization (Figure 4C). By quantifying the total bioluminescence flux in the entire mouse body, no differences between both infected groups were observed over time (Figure 4D). In order to further compare the kinetics of bacterial colonization of mice organs, we orally infected BALB/c mice with the WT or the Δ*dotB* strains, and all known infected organs were collected at days 5 or 7 pi and homogenized for CFUs counting: small intestinal tissue and its content, cæcum, colon tissue and its content, MLNs, spleen and liver (Figure 4E). Surprisingly, no statistical differences in CFUs were quantified at both investigated time points in all infected organs between WT- and Δ*dotB-*infected mice. These results indicate that, whereas the *y*T4BSS contributes to virulence in orally infected mice, it does not modulate organ colonization nor bacterial survival within these organs.

### The *y*T4BSS is associated to caecal and mesenteric lymph node necrosis

To investigate the mechanisms underlying the virulence defect upon oral infection of the Δ*dotB* mutant without being affected in organ colonization, we explored the tissue architecture by histological analyses of two key organs in the infectious process, the cæcum and the MLNs (Barnes et al., 2006; Fahlgren et al., 2009). BALB/c mice were infected with the WT or the Δ*dotB* mutant strains, organs were extracted at 7 dpi, fixed by immersion in 10% neutral-buffered formalin and processed for hematoxylin and eosin (HE) labeling. Multiple histological features were scored according to the degree of severity (0: absence, 1: mild, 2: moderate, 3: severe) in both organs (Table 6 and Materials and Methods), the resulting sum of scores provides an overall indication of the organ’s histopathology (Figure 5A and I). In both organs, the global scores were similar between WT or Δ*dotB* mutant infected mice. We may note in MLNs, global scores more spread between Δ*dotB*-infected mice than WT-infected mice, suggesting more heterogeneity regarding Δ*dotB* infection (Figure 5A and I). In MLNs, upon analysis of the recorded criteria separetely, the presence of multifocal to coalescing necrotizing granulomas was the most significant histological finding, showing more severe lesions in the WT-infected group than in the Δ*dotB*-infected group (Figure 5B and Figure S7). The severe lesions were caracterized by a central necrosis area with bacterial colonies and some degenerated neutrophils, surrounded by a rim of epithelioid macrophages (Figure 5C-D). Samples were thus subjected to histomorphometry analyses to measure the total percentage of necrosis area in each MLNs sample (Figure 5E and F). Interestingly, the percentage of necrosis area in MLNs of the WT-infected group was on average higher than in the case of the Δ*dotB*-infected group, the last composed of two distinct pattern: two mice highly affected (Figure 5G-H) and three mice with low necrosis areas. Thus, the *y*T4BSS might contribute to more tissular damage in MLNs, inducing more severe necrotizing granulomas and thus larger necrosis areas. In the cæcum, the presence of necrotic areas was also the most significant histopathological findings, with a higher severe score in the WT-infected group compared to the Δ*dotB*-infected group (Figure 5J-K, N-O and Figure S7). Histomorphometric analyses also revealed a higher average percentage of necrosis area in the cæcum of WT-infected mice than in the Δ*dotB*-infected mice (Figure 5L-M). Thus, the *y*T4BSS could be implicated in the development of necrosis in MLNs and in cæcum.

**Figure 5:**
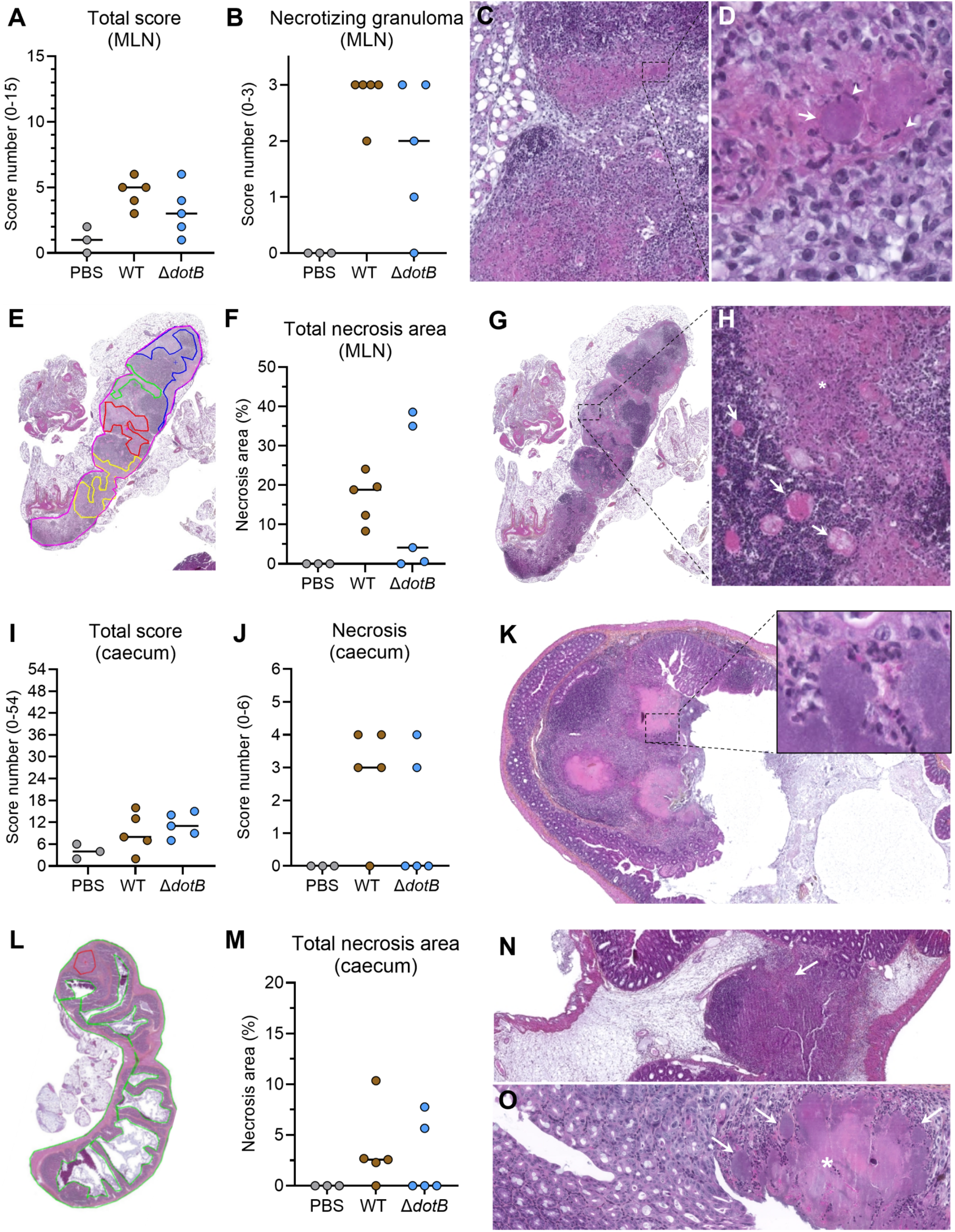
The *y*T4BSS is implicated in necrosis area formation in MLNs and cæca of infected mice. Mice were orally infected with WT or Δ*dotB* mutant strains (6*×*10^6^ CFU/mouse) or non-infected (PBS), cæca and mesenteric lymph nodes (MLNs) were collected at 7 dpi for histology analyses. **(A)** Total score calculated in MLNs on multiple histological findings with a maximal level of 15. **(B)** Severity level of the feature corresponding to necrotizing granuloma in MLNs. 0: absence of necrotizing granuloma, 1: mild lesion, 2: moderate lesion, 3: severe lesions. **(C)** Example of multifocal to coalescing necrotizing granulomas in MLNs from a WT-infected mouse (score 3, severe). **(D)** Higher magnification of the necrotizing granuloma. A central area of necrosis with the presence of bacterial colonies (arrow), degenerated neutrophils (arrowheads), and surrounded by a rim of epithelioid macrophages is evident. **(E)** Histomorphometric analysis of the percentage of necrosis in MLNs with Nikon NIS-Ar software. Yellow, blue, red and green colours in cæcum represent the necrotic areas measured. **(F)** Percentage of necrosis area on the total MLNs area. **(G)** Example of coalescing necrotizing granulomas in MLNs from a Δ*dotB*-infected mouse (score 3, severe). **(H)** Higher magnification of the necrotizing granulomas. The asterisk shows multiple degenerated neutrophils. Congestion of blood vessels is also evident (arrows). **(I)** Total score on multiple histological features measured in cæcum samples with a maximal level of 54 (listed in Table 6). **(J)** Severity level score of cæcum necrosis finding. 0: absence of necrosis to 6: high severity level. **(K)** Example of the necrotic area in the cæcum from a WT-infected mouse (score 3, severe). Inset shows the presence of bacteria and neutrophils. **(L)** Histomorphometric analysis of the percentage of necrosis in cæcum with Nikon NIS-Ar software. Red colour represents the necrotic area measured. **(M)** Percentage of necrosis area on the total cæcum area. **(N)** Example of the necrotic area (arrow) in the cæcum from a Δ*dotB*-infected mouse (score 2, moderate). **(O)** Example of the necrotic area (asterisk) in the cæcum from a WT-infected mouse (score 3, severe). Arrows show bacteria colonies.

We then conducted immunohistochemistry (IHC) analyses on neutrophils (anti-Ly6B2 antibody) and macrophages (anti-F4/80 antibody) in cæcal and MLNs samples. Samples were submitted to digital image analysis to calculate the percentage of positivity with respect to the whole area. Concerning Ly6B2 marker, in MLNs and cæca the percentage of positive Ly6B2 staining were similar between both infected groups (WT and Δ*dotB* mutant), with similar location of Ly6B2 positive cells (Figure 6A and E). In MLNs, positive Ly6B2 cells were detected surrounding the granulomas, outside of the epithelioid cells which form the lesions (Figure 6C-D). In cæca, neutrophils positive to Ly6B2 (and some macrophages-like cells also Ly6B2^+^) were clustered in caecal Peyer’s patches near to the necrotic areas (Figure 6F arrows and inset) and some were diffusely distributed in the lamina propria of the cæcal mucosa (Figure 6G).

**Figure 6:**
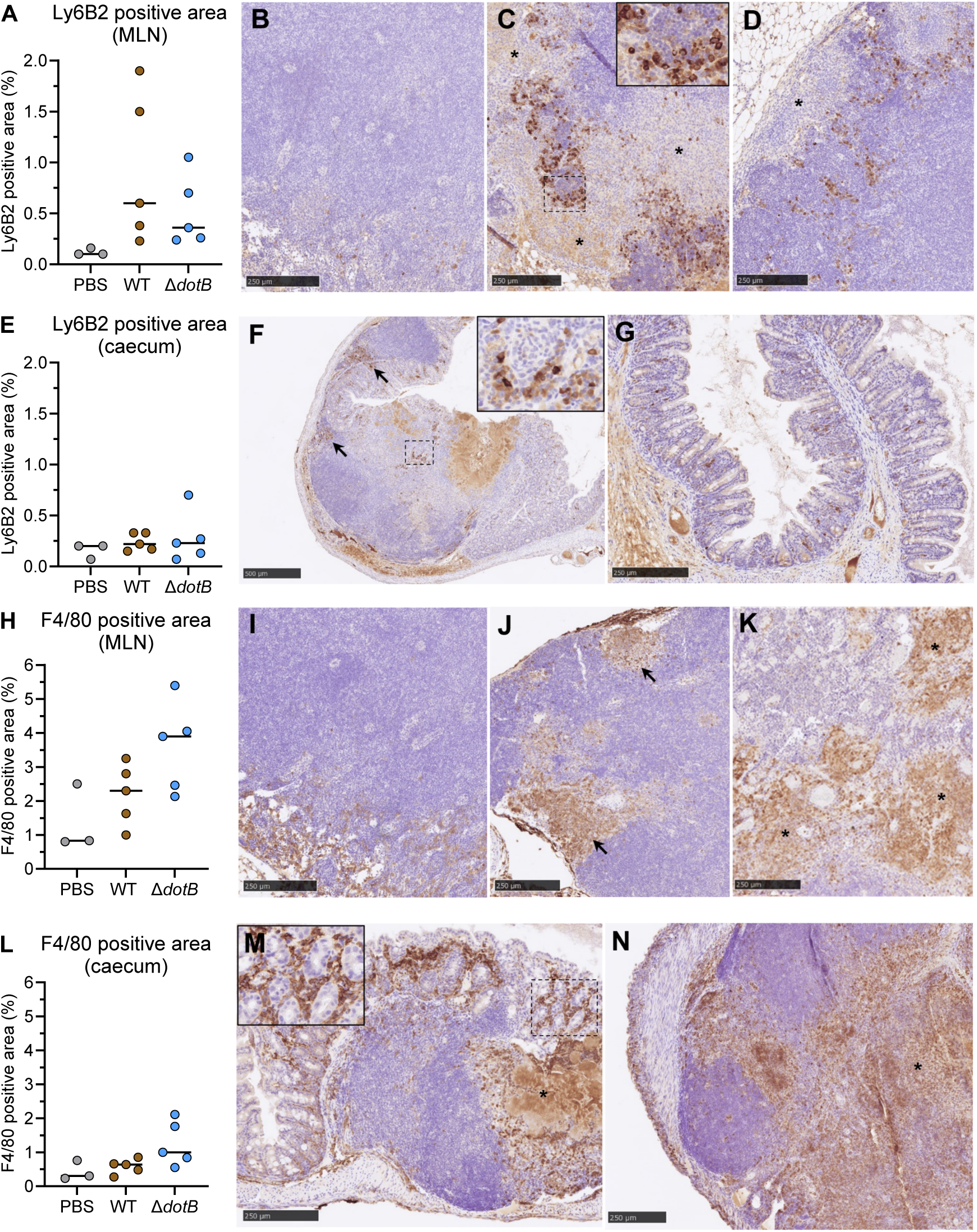
Immunohistochemistry analyses on neutrophils and macrophages in MLNs and cæcum of infected mice. **(A)** Percentage of positive area against Ly6B2 marker (neutrophils) in MLNs. **(B)** Representative picture of Ly6B2 in MLNs in non-infected mouse (control with PBS), **(C)** example in a WT-infected mouse. Inset shows detail of Ly6B2^+^ cells and **(D)** example in a Δ*dotB*-infected mouse. Asterisks show granulomatous lesions. **(E)** Percentage of positive area against Ly6B2 marker (neutrophils) in cæca. **(F)** Representative picture of Ly6B2 in cæcum in WT-infected mouse. Arrows and inset in show clusters of Ly6B2^+^ cells. **(G)** example in Δ*dotB*-infected mouse. **(H)** Percentage of positive area against F4/80 marker (macrophages) in MLNs. **(I)** Representative picture of F4/80 in MLNs in non-infected mouse (control with PBS). **J)** Example in a WT-infected mouse (arrows showing granulomas), and **(K)** Δ*dotB*-infected mouse (asterisks show necrotizing coalescing granulomas). **(L)** Percentage of positive area against F4/80 marker (macrophages) in cæca. **(M)** Representative pictures of F4/80 in cæcum in WT-infected mouse. Asteriks show F4/80^+^ cells in the periphery of a necrotic area in a Peyer’s patch. Inset shows detail of F4/80^+^ cells in the submucosa. **(N)** and Δ*dotB*- infected mouse. Asterisks show F4/80^+^ cells in the periphery of a necrotic area in a Peyer’s patch.

Interestingly, concerning the macrophage F4/80 marker, a tendancy for a higher average of positive stained areas were found in Δ*dotB*-infected mice than WT-infected mice in MLNs as well as in cæca (Figure 6H and L). Despite higher F4/80 positive areas in Δ*dotB*-infected mice, the localization of F4/80 positive cells were similar between WT- and Δ*dotB*-infected groups. In MLNs, the staining was detected for both groups (WT and Δ*dotB*) in the cytoplasm of macrophage-like cells, located in epithelioid cells of the granulomas (Figure 6J) and necrotizing granulomas (Figure 6K) as well as interspersed in the medulla of the lymph node and within the medullary cords. In the cæcum, F4/80 staining was detected in the cytoplasm of macrophage-like cells from the lamina propria of the caecal mucosa (Figure 6M), especially in the periphery of necrotic areas (Figure 6M and N). Thus, according with these results, it seems that in presence of the *y*T4BSS, less numbers of macrophages are present in cæca and MLNs from infected animals.

In conclusion, the *y*T4BSS might be implicated in the induction of necrosis in MLNs and the cæcum in infected mice, with a lower macrophagic inflammatory response present at these sites of infection.

### The *y*T4BSS modulates cytokine immune responses

Because macrophages appear to be a cell population targeted by the *y*T4BSS, we therefore sought to test whether the *y*T4BSS is associated with modulation of the cytokine responses to *Y. pseudotuberculosis* infection using mBMDM from BALB/c mice. For this, we measured the protein levels of a pannel of 18 cytokines and chemokines (Table 5) involved in various aspects of the immune response (innate and adaptative responses) in the supernatant of mBMDM upon 8 h of infection with the WT, the Δ*dotB* mutant or the complemented strain Δ*dotB::dotB.* Five cytokines: CCL2, G-CSF, IL-6, IL-33 and IL-10 displayed a higher concentration in the supernatant of mBMDM infected with the WT strain than in cells infected with the Δ*dotB* mutant (Figure 7, Figure S8). The complemented strain Δ*dotB::dotB* restored the WT phenotype in CCL2 and G-CSF cases but not statistically in IL-33, IL-6 and IL-10, even if the tendancy followed the one of the WT. Thus, the *y*T4BSS is associated with an increase in cytokine production from the innate immune response of mBMDM upon *Y. pseudotuberculosis* infection.

**Figure 7:**
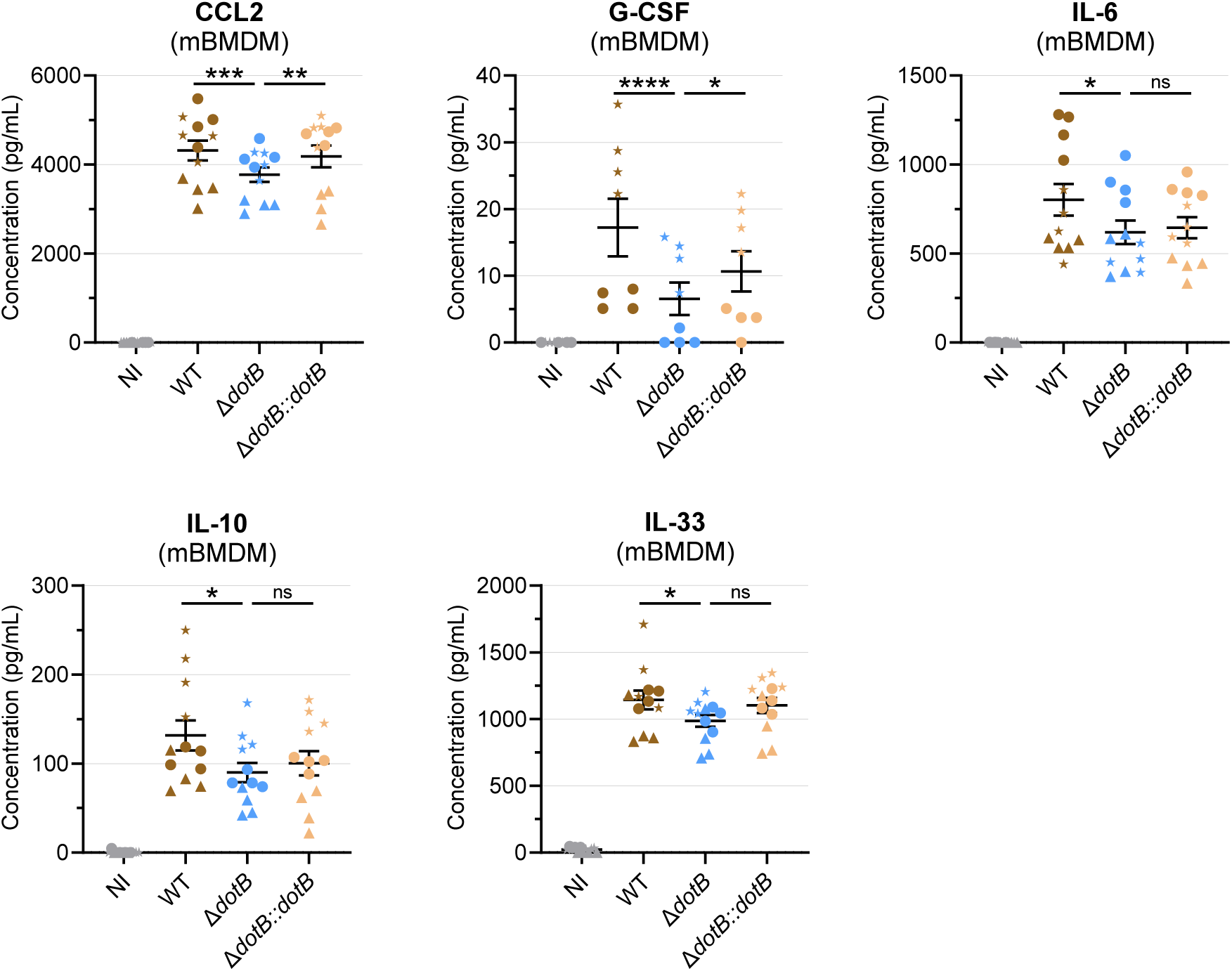
The *y*T4BSS modulates the cytokine immune response. Cytokine profile analysis on supernatants from infected mBMDMs at 8 hpi with WT, Δ*dotB* mutant or Δ*dotB::dotB* complemented strains (MOI 10) using the Luminex technology. Data are representative of 3 independent experiments (triangle, round and stars) with 4 technical replicates and presented as mean and SEM. Mixed model with Tukey’s multiple comparisons correction were performed on logY+1 transformed data.

## Discussion

*Y. pseudotuberculosis* is an enteric pathogen, traditionnally associated to mesenteric adenitis in humans, worldwide. A phylogenetic cluster of strains from lineages 1 and 8 have been implicated in the undercharacterized syndrome FESLF, causing specific clinical manifestations of skin rash, hyperemic tongue and desquamation, with cases still reported mainly in Russia, Korea and Japan (Ocho et al., 2018; Sato, 1987; Suzuki et al., 2023). For the time being, to the best of our knowledge, the only virulence factor linked to FESLF is the chromosomally encoded superantigen YPMa, whose presence is correlated with systemic symptoms found in FESLF patients (Abe et al., 1997; Abe et al., 2003; Ueshiba et al., 1998). The plasmid pVM82, present in FESLF strains from lineage 8, had been previously associated to bacterial pathogenicity but no specific virulence factors have been characterized so far in this genetic element (Dubrovina et al., 1999; Gintsburg et al., 1988; Sever et al., 1991). In the continuity of the work of Eppinger et al., 2007 who identified in the pVM82 a T4BSS harboring *dot/icm* genes, we investigated in this work the specificities of the *y*T4BSS as well as its contribution to virulence.

By analyzing the structure of the *y*T4BSS gene cluster, we identified three new genes: *icmT* within the previous identified gene cluster, as well as two copies of *icmX* (*icmX1* and *icmX2*) in a distant gene cluster on the pVM82. By comparing gene cluster homologies with known functional T4BSS systems, we found that *y*T4BSS genes display higher homology with a greater number of genes with those of the recently described T4BSS of *P. putida* than with those of *L. pneumophila* and *C. burnetii*. The only genes missing in *Y. pseudotuberculosis* compared to the closest strain *P. putida* are *icmW* and *dotU/icmH*. In *L. pneumophila*, IcmW forms a stable chaperone protein complex with IcmS allowing translocation of IcmSW-dependent effectors (Ninio et al., 2005). Because *icmS* is also absent in the *y*T4BSS, we could hypothesize that effector translocation is only DotM-dependent or that other chaperone systems still need to be deciphered in *Y. pseudotuberculosis*. In *L. pneumophila*, DotU is dependent on IcmF, both integral inner membrane proteins, for T4BSS polar assembly and structural stability of the DotH, DotG and DotF subunits (Ghosal et al., 2019; Sexton et al., 2004). Because the *y*T4BSS lacks *icmF* and *dotU*, we could hypothesize that the *y*T4BSS might display structural specificities, different from the well-defined structure of the T4BSS in *L. pneumophila*. Thus, the *dot/icm* genes set of *Y. pseudotuberculosis* SP-1303 appears to be complete, encoding main structural and enzymatic components of a typical T4BSS, making it a potentially functional system.

By testing different *in vitro*, *in cellulo* and *in vivo* conditions, we found that *y*T4BSS genes are continuously expressed in rich culture media and also during *Y. pseudotuberculosis* intracellular life cycle, as well as during mouse infection without organ specificity. One key parameter found to influence the rate of expression appears to be temperature, with higher expression rate encountered at higher temperature (37°C) than lower temperature (21°C, 28°C), suggesting for a role of the *y*T4BSS in the mammalian host. In comparison to other T4BSS models, high temperatures do not seem to be an inductive signal for *dot/icm* gene expression. Indeed, within phagocytes, *dot/icm* and related effectors expression is induced by nutrient deficiency in the *Legionella*-containing vacuole (LCV), whereas acidic pH and specific amino acid composition (phenylalanin, proline and serine) are signals inducing *dot/icm* gene expression in the *Coxiella*-containing vacuole (CCV) (Jaboulay et al., 2021; Al-Khodor et al., 2009; Newton et al., 2020). Concerning *P. putida*, no gene expression study has been performed so far, but the T4BSS is functional in bacterial killing in environmental conditions (30°C) (Purtschert-Montenegro et al., 2022). Thus, the *y*T4BSS appears to be unique among T4BSS in terms of condition of expression, with upregulation found at 37°C. Other environmental conditions have to be explored (pH, growth phase, media composition…) in order to fully decipher conditions involved in *y*T4BSS gene regulation. In *Y. pseudotuberculosis*, many other virulence factors are also upregulated at 37°C, such as the T3SS and Yops, the majority of adhesins (YadA, Ail, PsaA), a type IV pili, making the *y*T4BSS an additional molecular weapon in the context of host infection.

To our surprise, even if the *y*T4BSS is expressed through the intracellular life cycle of *Y. pseudotuberculosis*, it does not take part in bacterial survival, nor in the biogenesis of YCVs in epithelial cells and macrophages. Thus, the *y*T4BSS does not participate in cellular infection in *Y. pseudotuberculosis*, as opposed to the T4BSS of the obligate intracellular pathogens *L. pneumophila* and *C. burnetii*, which are totally dependent on their T4BSS for their intracellular life cycle in phagocytes (Qiu and Luo, 2017). Using fluorescent microscopy and transmission electron microscopy, we described the biogenesis of a single-membrane vacuole in HeLa cells, containing replicative *Y. pseudotuberculosis,* which increases in size over time, localizes close the host nucleus, as it was also shown by Ligeon et al, independently of *y*T4BSS functions (Ligeon et al., 2014). To date, because no bacterial determinants have been linked to YCV biogenesis, the use of a mutant library of *Y. pseudotuberculosis* isolates coupled to cellular infection analyses by microscopic screening would allow to fill this gap in the literature.

We found a key role of the *y*T4BSS in *Y. pseudotuberculosis* FESLF strain pathogenicity during oral infection of mice but not during intravenous inoculation, demonstrating that the *y*T4BSS is a new virulence factor, whose function depends on the infection route. Interestingly, the superantigen YPMa from FESLF strains also acts as a virulence factor depending on the infection route. Unlike the *y*T4BSS, YPMa plays an important role in virulence during intravenous infection whereas it has no effect during intragastric infection (Carnoy et al., 2000). Nevertheless, the *y*T4BSS does not take part in bacterial colonization of organs or survival during oral infection, as demonstrated by similar bacterial burden between the WT strain and the Δ*dotB* mutant strain, which was also found to be similar between WT and Δ*ypmA* mutant strain (Carnoy et al., 2000). Similar observations have been made in *Staphylococcus aureus* in which the nicotinamide adenine dinucleotide kinase (NADK) is essential for full virulence in a model of zebrafish infection leading to neutropenia and death, whereas NADK absence does not impact bacterial burden in fish, while it leads to greater fish survival (Leseigneur et al., 2022). Performing histological analyses on key organs during the infectious process, cæcum and MLNs, we observed the involvement of the *y*T4BSS in induction of high severity lesions corresponding to important necrosis areas in the cæcum and in MLNs at sites of necrotizing granulomas. For the time being, we do not know if these necrotic lesions are the direct effect of *y*T4BSS functions on host cells or indirect consequences of *y*T4BSS functions. Necrosis has already been described in pyogranuloma in MLNs and along the gastrointestinal tract upon infection with *Y. pseudotuberculosis* lacking the *y*T4BSS (El-Maraghi and Mair, 1979; Sorobetea et al., 2023). Thus, the cell composition of necrotic areas and dynamic of necrosis formation, depending on the *y*T4BSS, has to be deciphered. We additionally observed in a *y*T4BSS-dependent manner, a tendency for a lower percentage of macrophages found in infected MLNs and the cæcum. One hypothesis linking important necrosis areas with a reduced macrophage population could be that macrophages are a cell population targeted for cell death by the *y*T4BSS, participating in appearance of large necrotic lesions. However, the cellular targets and the type of cell death involved remain to be identified.

Thanks to cytokine profile analyses, we observed that the *y*T4BSS appears to participate in modulation of immune responses using a model of mBMDM infection *in vitro*, leading to higher levels of innate immunity related cytokines and chemokines production. *L. pneumophila,* thanks to its T4BSS, also modulates the host immune response, inducing an increased pro-inflammatory cytokine response in a mBMDM infection model as well as in an intranasal mouse model, resulting in IL-1α and IL-1β production, which participate in neutrophil recruitment to the lungs of infected animals (Barry et al., 2013; Copenhaver et al., 2015). However, *C. burnetii* has opposite effect by silencing host innate immune responses in a T4BSS-dependent manner by downregulating NF-κB signaling pathways (Burette et al., 2020; Mahapatra et al., 2016). Thus, the *y*T4BSS shares common functions with the T4BSS of *L. pneumophila* in increasing inflammatory cytokines responses, although the precise molecular mechanisms remain to be identified in *Y. pseudotuberculosis* FESLF strains. *Y. pseudotuberculosis* also depends on its T3SS for infection, which displays immunosuppressive properties, with YopJ for example known to inhibit pro-inflammatory cytokines production (such as IL-6, IL-8, TNF-α) in macrophages and epithelial cells, or YopM inhibiting production of IL-1β and TNF-α, but increasing IL-10 production in macrophages (Berneking et al., 2023; Kerschen et al., 2004). Because of opposite impacts of *y*T4BSS and T3SS on the host cytokines responses, the interplay between both systems needs to be explored. We could hypothesize that depending on the phase of the disease, organ or cell type, *y*T4BSS and T3SS together or sequentially, impact the host immune system, participating in *Y. pseudotuberculosis* pathogenicity. Sequential activation of secretion systems is encountered in the example of *Salmonella enterica* serovar typhimurium, starting with the T3SS from the *Salmonella* pathogenicity island 1 (SPI-1) allowing invasion of intestinal epithelial cells and induction of inflammatory responses, then within macrophages T3SS from SPI-2 is activated and in later phases of infection, its T6SS participates in persistent infection (Das et al., 2013). In *Y. pseudotuberculosis* FESLF strains, the superantigen YPMa has also an impact on the host immune system but acts on the adaptive immune response by binding specific CD4^+^ and CD8^+^ T cells, leading to their activation and expansion with high production of IL-2 and IL-4 (Goubard et al., 2015; Miyoshi-Akiyama et al., 1995; Seprényi et al., 1999; Uchiyama et al., 1993). In our study, IL-2 and IL-4 levels are not influenced by the presence of the *y*T4BSS. Thus, with an impact only during oral infection and on inflammation, the *y*T4BSS appears to participate in the first steps of infection, whereas with a virulence role only during intravenous infection and on T cell populations, YPMa acts later in the infection process, probably when bacteria reach the bloodstream, as *ypmA* expression is induced in liver and spleen (Goubard et al., 2015).

The *y*T4BSS appears to have an impact on mammalian host at the immune response level. However, *y*T4BSS genes are highly homologous to those of the plant root colonizer *P. putida*, which functions as a contact-dependent bacterial killing machinery, targeting a wide range of soil and plant-associated Gram-negative bacteria, allowing *P. putida* to persist in its environment (Purtschert-Montenegro et al., 2022). As already described in *Vibrionaceae* family with a T6SS harboring antibacterial and anti-eukaryotic activities, the potential properties in bacterial killing of the *y*T4BSS remain to be explored (Dar et al., 2018).

To conclude on this study, we demonstrated the involvement of a new virulence factor, a *y*T4BSS, in the pathogenicity of *Y. pseudotuberculosis* strains from lineage 8 responsible for FESLF. Although the system is not involved in the intracellular life cycle of *Y. pseudotuberculsosis*, the *y*T4BSS is highly expressed under mammalian host temperatures, participates in induction of necrotic lesions in infected tissues and modulates host cytokine immune responses. Further studies characterizing the effectors secreted by the *y*T4BSS should allow to better understand its contribution to *Y. pseudotuberculosis* infection.

## Materials and Methods

### Bacterial strains

Bacterial strains and plasmids used in this study are listed in Table 1 and Table 2. *Y. pseudotuberculosis* and *E. coli* strains were grown in lysogeny broth (LB) medium (Difco) under stirring or on LB agar plates at 28°C for 48 h or 37°C for 24 h respectively. When required, culture medium was supplemented with antibiotics (irgasan 0.1 µg/mL carbenicillin 100 µg/mL, kanamycin 30 µg/mL, chloramphenicol 25 µg/mL).

**Table 1:**
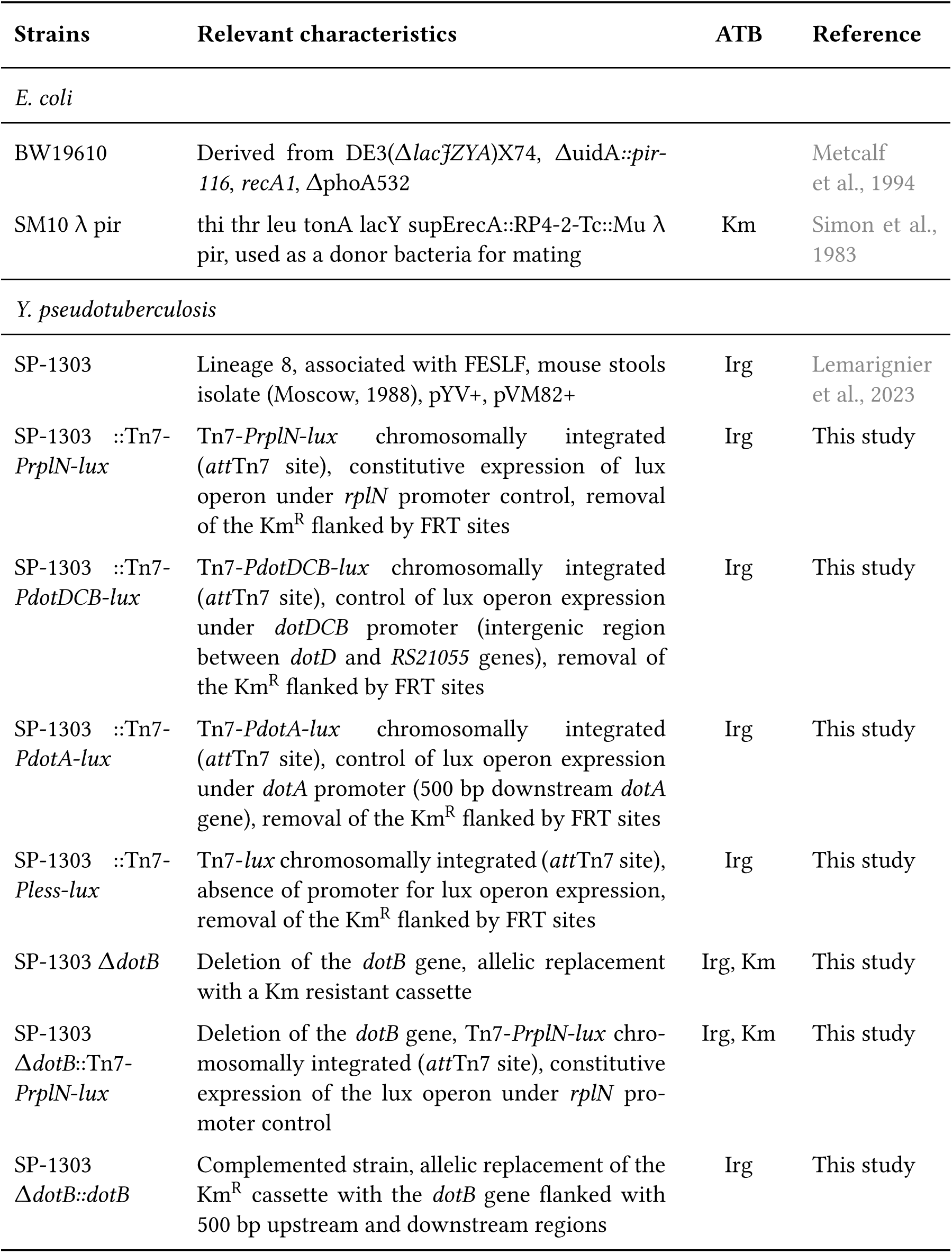
Stain used in this chapter. ATB: Antibiotic resistance.

**Table 2:** Plasmids used in this chapter. ATB: Antibiotic resistance.

| Plasmids | Relevant characteristics | ATB | Reference |
| --- | --- | --- | --- |
| pTNS2 | Helper suicide plasmid encoding the specific TnsABC+D transposition pathway for mini Tn7 transposition | Amp | Choi et al., 2005 |
| pFLP3 | Flp recombinase-encoding plasmid, β-lactamase encoding gene | Amp | Choi et al., 2005 |
| pUC18R6K mini-Tn7-*Km<sup>R</sup>-Pless-luxCDABE* | Suicide mini-Tn7-*Km<sup>R</sup>-lux* delivery vector, absence of promoter upstream of the lux operon | Km | Derbise et al., 2019 |
| pUC18R6K mini-Tn7-*Km<sup>R</sup>-PrpLN-luxCDABE* | Suicide mini-Tn7-*Km<sup>R</sup>-PrpLN-lux* delivery vector | Km | Derbise et al., 2019 |
| pUC18R6K mini-Tn7-*Km<sup>R</sup>-PdotDCB-luxCDABE* | Suicide mini-Tn7-*Km<sup>R</sup>-PdotDCB-lux* delivery vector (*dotDCB* promoter cloned into the SpeI/XmaI sites of pUC18R6K-Tn7-*Km<sup>R</sup>-Pless-luxCDABE*) | Km | This study |
| pUC18R6K mini-Tn7-*Km<sup>R</sup>-PdotA-luxCDABE* | Suicide mini-Tn7-*Km<sup>R</sup>-PdotA-lux* delivery vector (*dotA* promoter cloned into the SpeI/XmaI sites of pUC18R6K-Tn7-*Km<sup>R</sup>-Pless-luxCDABE*) | Km | This study |
| pUC4K | Kanamycin resistant cassette amplified with primers Km-F/Km-R | Km | Taylor and Rose, 1988 |
| pKOBEG-sacB | $\lambda$ phage red $\gamma$ $\beta$ $\alpha$ operon expressed under the control of the arabinose-inducible pBAD promoter, *sacB* cloned at NdeI restriction site | Cm | Derbise et al., 2003 |
| pCVD442 | *sacB*, β-lactamase encoding genes | Amp | Donnenberg and Kaper, 1991 |
| pCVD442-*up-dotB-down* | Insertion at SalI and SphI sites of the entire region containing *dotB* gene flanked with 500 bp of upstream and downstream regions | Amp | This study |

### Search for T4BSS genes, oriT and relaxase

The presence of T4SS genes and relaxase was assessed on multiple pVM82 sequences from *Y. pseudotuberculosis* FESLF strains using MacsyFinder v2.07rc (Néron et al., 2023) with the “Plasmids” models of the CONJScan module. Briefly, MacSyFinder uses HMM protein profiles and a set of rules (which are defined in the models) about their presence and proximity. Macsyfinder was used with default parameters and CONJScan models were ran all at once with the parametters “- all”. A complementary approach was used with oriTfinder, allowing to search for an origin of transfer (oriT) on the pVM82 plasmid (Li et al., 2018).

### Phylogenetic reconstruction and presence of virulence factors

In a total of 287 genomes of *Y. pseudotuberculosis* (including 20 *Y. pestis*), the nucleotidic sequences of the 500 genes of the cgMLST-*Yersinia* (Savin et al., 2019) were extracted and aligned using mafft v7.467 (Katoh and Standley, 2013). From this alignment, a phylogenetic reconstruction was performed with IQ-TREE2 (Minh et al., 2020) using a bootstrap value of 1,000 and by first determining the best substitution model thanks to the “ModelFinder” implemented in IQ-TREE2. ABRIcate v.1.0.1 (github.com/tseemann/abricate) was used to search for the presence of the different coding sequences of the *y*T4BSS together with the different versions of the *ypm* gene in the 2,015 genomes. For this purpose, an in-house database containing the sequences of the 18 genes was created for ABRIcate. Default parameters of 80% identity and 80% coverage were used to identify the presence of the genes. Visualization of the phylogenetic reconstruction together with the presence of the virulence genes was generated using phandango (Hadfield et al., 2018). The genotypes of *Y. pseudotuberculosis* were plotted on the tree as described in (Savin et al., 2019).

### *y*T4BSS gene cluster conservation comparison

Homologous gene cluster searches for the *y*T4BSS operon of the strain *Y. pseudotubeculosis* SP-1303 (GenBank accession number: NZ_CP130902.1) was performed on the *Yersinia* spp database of NCBI by using cblaster pipeline in default mode (minimum identity 30%, minimum query coverage 50%). NCBI identifiers used were: WLF06076.1 (IcmX1), WLF06077.1 (IcmX2), WLF06128.1 (DotA), WLF06119.1 (DotM/IcmP), WLF06113.1 (DotB), WLF06112.1 (DotC), WLF06111.1 (DotD), WLF06108.1 (DotI/IcmL), WLF06107.1 (DotH/IcmK), WLF06106.1 (DotG/IcmE), WLF06105.1 (DotF/IcmG), WLF06104.1 (DotE/ IcmC), WLF06102.1 (DotN/IcmJ), WLF06100.1 (DotO/IcmB), WLF06099.1 (DotL/IcmO), as well as amino acid sequence of IcmT newly described (Gilchrist et al., 2021). The gene cluster conservation comparison of the *y*T4BSS of *Y. pseudotuberculosis* SP-1303 and other T4BSS members was performed by using clinker/0.0.23 pipeline with default parameters (minimum alignement sequence identity of 0.3) by giving Genbank files of the selected regions of the IncI plasmid R64 (NC_005014) and of the following strains: *Coxiella burnetii* RSA 493 (NC_002971), *Legionella pneumophila* Philadelphia 1 (NC_002942), *Pseudomonas putida* IsoF (CP072013.1). Clinker performs pairwise local or global alignments between every sequence in every unique pair of clusters (Gilchrist and Chooi, 2021).

### Synteny analysis and representation

The IP31758 strain was used as reference strain and aligned with 45 strains selected for their association with FESLF and presence of the pYV plasmid. Analysis was performed as described in (Lê-Bury et al., 2023). Briefly, a protein BLAST search was automatically performed for each gene of each genome against the other genomes to construct the homolog database, using the BLAST1 command-line tool v2.13.0 (Camacho et al., 2009). Synteny was constructed using a best-hit bidirectional BLAST search and implemented in the SynTView framework (Lechat et al., 2013). The tool was reprogrammed from Flash to JavaScript with the HTML5 2d Canvas JavaScript library Konva (konvajs.org), which enables high-performance animations. Drawing strategies were developed to ensure that graphical outputs remain below the technological limits of drawing more than 10,000 dynamic graphical objects, using a mix of static and dynamic objects. SynTViewJS can be used on a public web application at plechat.pages.pasteur.fr. The code repository is at gitlab.pasteur.fr/plechat/syntviewjs.

### Generation of a **Δ***dotB* mutant and complemented strains

SP-1303 Δ*dotB* mutant was generated by allelic replacement of *dotB* (*dotB*_RS21065 from start to stop codons) by homologous recombination with a linear three-step PCR fragment containing a kanamycin resistant cassette, flanked with 500 bp upstream and downstream regions of *dotB*. The detailed method followed is described in (Derbise et al., 2003). Primers used are listed in Table 3. The SP-1303 Δ*dotB::dotB* complemented strain was generated by allelic replacement of the kanamycin resistant cassette of the Δ*dotB* mutant with the original *dotB* gene cloned in the pCVD442 plasmid. For that, the entire dotB sequence with 500 bp flanked regions was PCR amplified from a SP-1303 genomic DNA extract using a high-fidelity polymerase (Phusion, NEB). Purified amplicons were cloned at SalI and SphI sites (NEB enzymes) into the pCVD442 plasmid and electroporated in *E. coli* SM10 donor bacteria. Mating was performed between *E. coli* SM10 pCVD442-*up*-*dotB*-*down* and the SP-1303 Δ*dotB* mutant by mixing equivalent volumes of adjusted overnight (O/N) liquid culture to OD=2 on a 0.45 µm filter (peel corner, MERCK) positioned on a LB agar plate for 6 h at 28°C. After mating, bacteria were collected, serially diluted, and plated on LB-Irg-Km-Amp agar for 48 h. Resistant clones were cultured O/N in liquid LB and spread on LB agar 10% sucrose without NaCl for 48 h at 28°C, for resolution of the meroidiploid previously formed. The complemented strain was checked for its sensitivity to kanamycin and ampicillin. The complete genome of the Δ*dotB* mutant and its complemented strainΔ*dotB::dotB* were extracted using the PureLink Genomic DNA kit (Invitrogen) and checked for mutations using two complementary sequencing technologies: Oxford Nanopore Libraires (long reads) and Illumina (short reads). Librairies were prepared with the Nextera kit for Illumina NextSeq 550 sequencing by the Plateforme de microbiologie mutualisée (P2M) of Institut Pasteur and by using the Rapid Barcoding kit (SQK-RBK004) for R9.4.2 ONT MinION flow cell sequencing.

**Table 3:**
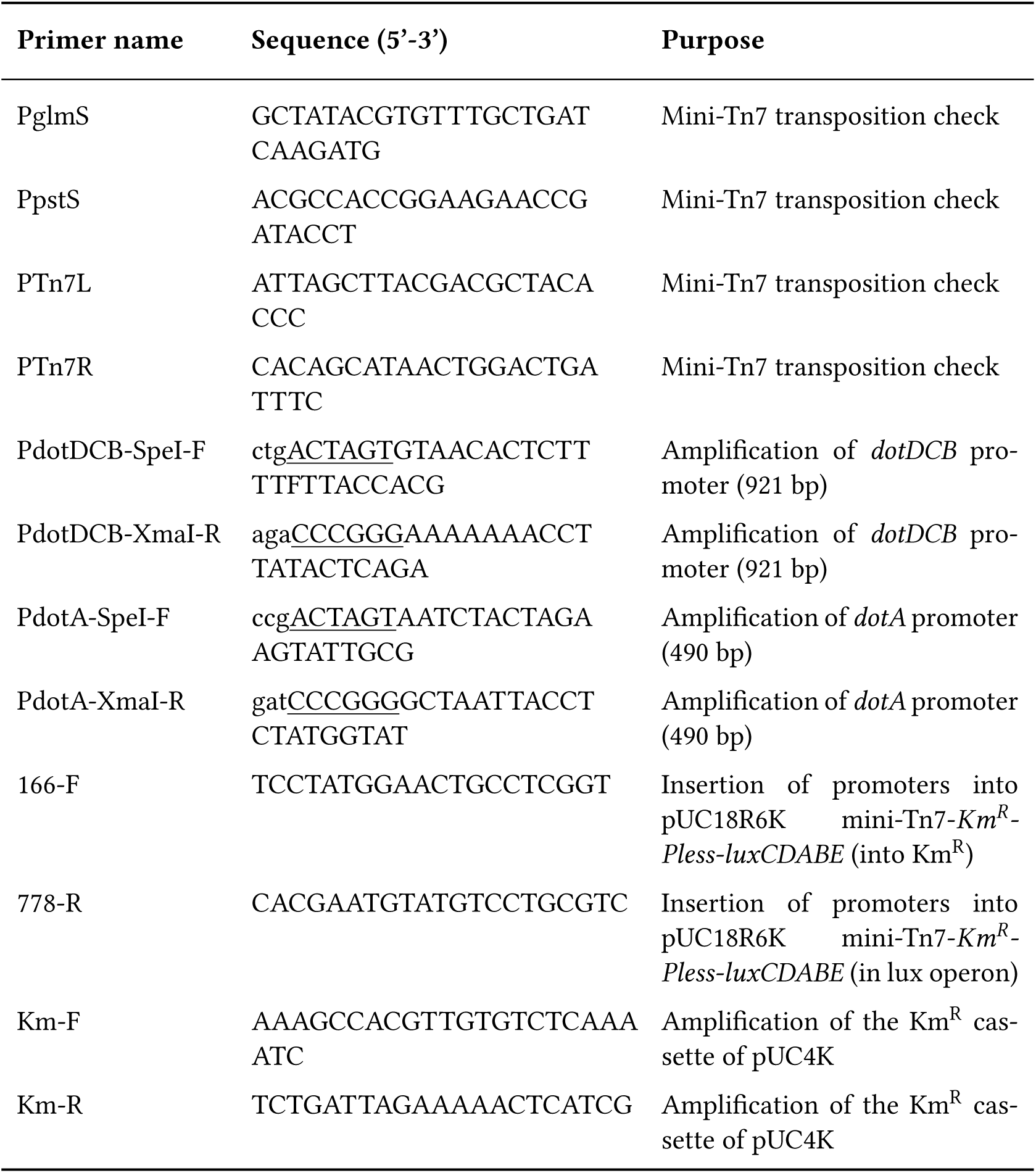
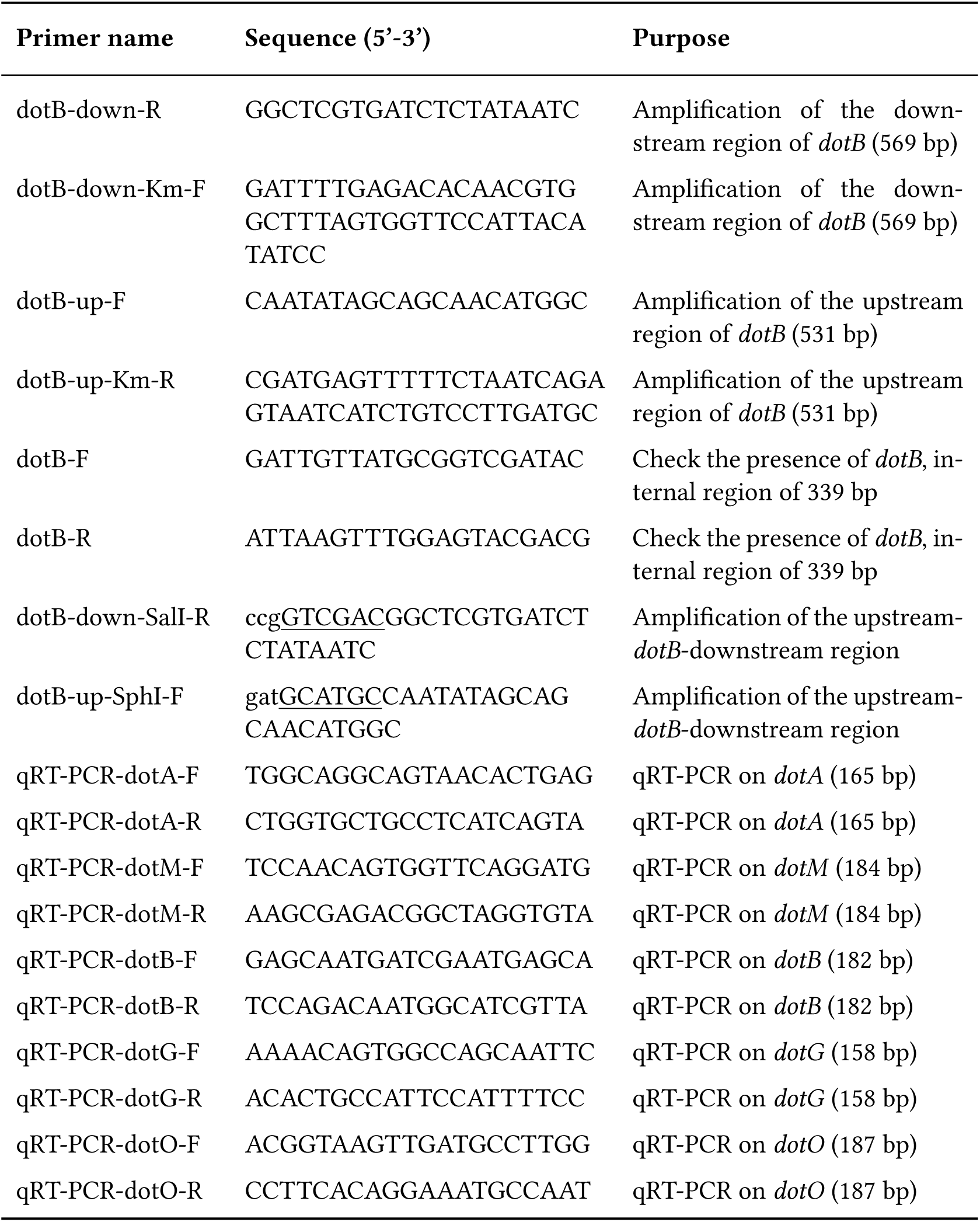
Primers used in this chapter. Underlined nucleotides indicate restriction sites used for cloning.

### Generation of bioluminescent reporters

Constitutively bioluminescent strains: SP-1303::Tn7-*PrplN-lux* and SP-1303 Δ*dotB*::Tn7-*PrplN-lux* were generated by insertion into the unique chromosomic *att*Tn7 site of the *luxCDABE* operon from *Photorabdus luminescens*, under the control of the constitutive promoter of *rplN* gene, by co-electroporated plasmids pUC18R6KT-mini-Tn7-*Km^R^*-*PrplN-luxCDABE* and pTNS2 as described in (Derbise et al., 2019). For the generation of transcriptional bioluminescent reporters, the promoter regions of *dotA* (490 bp upstream *dotA_RS21140* and *dotDCB* (intergenic region of 921 bp between *dotD_RS21055* and *RS21050* genes) were PCR-amplified using a high-fidelity polymerase (Phusion, NEB) and PCR products were purified before cloning (QIAquick PCR purification kit, QIAGEN). Amplicons were cloned in front of the *luxCDABE* operon in the plasmid pUC18R6KT-mini-Tn7-*Km^R^-luxCDABE* into the SpeI and XmaI restriction sites (highfidelity enzymes, NEB). Constructs were verified for mutations by sequencing, using primers listed in Table 3. The kanamycin resistant cassette flanked by FRT regions was removed in all constructions by using a recombinase-encoding plasmid pFLP3, which was then cured as detailed in (Derbise et al., 2019).

### Bioluminescence monitoring

#### In vitro

Reporter strains were grown O/N at 28°C and diluted in LB (50 mL) to reach OD_600nm_= 0.1 prior to culture at a tested temperatures : 21°C, 28°C, 37°C and 42°C. OD_600nm_ and photon emission were measured every 2 h, using a spectrophotometer and a Glomax Discover microplate reader (Promega) respectively as described in (Derbise et al., 2019). Photon emission in relative light units (RLU) were corrected by OD_600nm_ measures.

#### In cellulo

HeLa cells were seeded the day before infection at 10^4^ cells per well in 96-well plates. Reporter strains grown O/N at 28°C were adjusted to a multiplicity of infection (MOI) of 70 in cell culture media. Infected plates were centrifuged (188 g x 5 min) to synchronize bacterial invasion and were incubated for 1h before gentamicin addition (20 µg/mL) for the remaining time: 1h, 3h, 7h or 23h. Cell culture media was replaced by PBS prior to photon emission measurement using a Glomax Discover microplate reader (Promega). Cells were lysed in distilled water for intracellular bacterial counts on LB agar as described in (Kühbacher et al., 2013). Photon mesurements in RLU were corrected by CFUs counts.

#### In vivo

Real-time bioluminescence images were acquired on anesthetized mice (constant flow of 2.5% isoflurane mixed with oxygen) by using the In Vivo Imaging System (IVIS 100; Caliper Life Sciences) in automatic mode. To analyse the bioluminescence signal within organs, mice were euthanized and the gastrointestinal tract, pool of MLNs and spleen were isolated. Image analysis was performed using the Living Image 4.5 software (Revvity). Photon emission is comparable between mice by applying a common scale. Total photon emission was quantified by drawing regions of interest (ROIs) around the entire body and given as average radiance (photons/s/cm^2^/steradian [sr]).

### RNA extraction and quantitative Real-Time PCR

Bacteria were grown at 21°C, 28°C or 37°C in triplicate in LB under stirring conditions until late exponential phase (OD_600_=1.3), pelleted and frozen at −80°C. Pellets were resuspended in TRIzol (Invitrogen) at 4°C and processed for lysis in beads-containing tubes (#P000914-LYSK06A, Bertin Instruments) using a cooling homogenizer for two cycles of 30 s (Precellys24 and Cryolys, Bertin Instruments) and centrifuged for 15 min at 12,000*×*g at 4°C. The aqueous phase was mixed twice with chloroform, left for 5 min at room temperature and centrifuged. The clear aqueous phase was mixed with isopropanol and left for 5 min at room temperature for RNA precipitation. After centrifugation, the pellet was washed in 70% ethanol, left still for 5 min and transferred to a RNeasy spin column (RNeasy Mini kit, QIAGEN). The column was washed two times with RPE buffer before elution in RNA-free water. RNA samples were treated with DNase according to the manufacturer’s recommendations (TURBO DNase-free kit, Ambion) and quantified using a Qubit 4 Flurometer (Invitrogen). A quality control step on 1% agarose gel with guanidine thiocyanate and ethidium bromide was done. Retrotranscription was performed thanks to the iScript cDNA synthesis kit following the manufacturer’s instructions (Bio-Rad). Quantitative RT-PCRs (qRT-PCRs) were carried out in triplicate with three technical replicates with the SsoAdvanced Universal SYBR Green kit (Bio-Rad) on a CFX384 Touch Real-Time PCR Detection System (BioRad). The qRT-PCR program used was 98°C for 3 min, 40 cycles of 98°C for 15 s and 60°C for 30 s, then 95°C for 15 s, finishing with a melt curve from 60°C to 95°C with incrementation of 0.5°C. Primers used are listed in Table 3. Reference genes *nadB*, *proC* and *rpoD* were selected according to the following study (Koch et al., 2019). Data were analyzed with the ΔCt method (Livak and Schmittgen, 2001).

### Cell culture

HeLa cells, Caco-2 cells and RAW 264.7 (TIB-7) macrophages (ATCC) were cultured and maintained in DMEM supplemented with high glucose (4.5 g/L, Gibco) and 10% fetal bovine serum (FBS, Gibco). Mouse bone marrow-derived macrophages (mBMDM) or human blood monocyte-derived macrophages were grown in RPMI, high glucose (Gibco) with 10% FBS and 10mM Hepes. Addition of 50 µM β-mercaptoethanol and antibiotics (Penicilline/Streptomycin, Gibco) was used for mBMDM and humand blood monocytes-DM differentiation steps. Cells were incubated at 37°C, 5% O_2_.

### Human blood monocyte-derived macrophages isolation

Isolation of human blood monocyte-DM were performed as described in (Klunk et al., 2022). Briefly, blood mononuclear cells were isolated by density gradient centrifugation from buffy coats of human healthy donors (Ficoll-Paque Premium, Sigma Aldrich, St. Louis, MI, USA). Monocytes were purified from peripheral blood mononuclear cells by positive selection with magnetic CD14 MicroBeads (Miltenyi Biotech, Bergisch Gladbach, Germany) using the autoMACS Pro Separator. The purity of the isolated monocytes was verified using an antibody against CD14 (BD Biosciences) and only samples showing >90% purity were used to differentiate into macrophages. Monocytes were then cultured for 7 days in RPMI media supplemented with 10% FBS, L-glutamine (Fisher) and M-CSF (20ng/mL; R&D systems). Cell cultures were fed every 2 days with complete medium supplemented with the cytokines previously mentioned. Before infection, the differentiation/activation status of the monocyte-derived macrophages was checked by flow cytometry, using antibodies against CD1a, CD14, CD83, and HLA-DR (BD Biosciences). All samples presented the expected phenotype for non-activated macrophages (CD1a+, CD14+, CD83, and HLA-DRlow). human blood monocyte-DM were seeded at a concentration of 10^5^ cells/mL the day before the infection in media without M-CSF and antibiotics.

### Mouse bone marrow-derived macrophages (mBMDM) isolation

Tibias and femurs were dissected from euthanized 8-weeks old BALB/cByJ mice. Collection of bone marrow cells was performed by flushing cell culture media into the bone using a sterile syringe with a 25G needle. Harvested cells were centrifuged at 300*×*g for 5 min at 4°C and resuspended in red blood cell lysis buffer (#130-094-183, Miltenyi Biotec) for 5 min. Cells were centrifuged and resuspended in RPMI, high glucose (4.5 g/L, Gibco) supplemented with 10% FBS (Gibco) with antibiotics (Penicilline/Streptomycin Gibco) and seeded at 1*×*10^6^ cells/mL in a T150 cm^2^ flask (Corning) for incubation at 37°C, 5% O_2_. After 4h of incubation, non adherent cells were harvested and removed from attached epithelial cells. Collected cells were seeded at 1*×*10^6^ cells/mL in petri dish (Greiner Bio-one) in cell culture media supplemented with M-CSF (25 µg/mL, mouse M-CSF #130-101-704, Miltenyi Biotec) and antibiotics, and cultured at 37°C, 5% O_2_ for 5 days. Half of the media was changed and replaced by fresh culture media with M-CSF and antibiotics for two additional days of differentiation. mBMDM were seeded at a concentration of 5 x 10^5^ cells/mL the day before the infection in media without M-CSF and antibiotics.

### Gentamicin protection assays

The day before the experiment, cells were seeded at 10^5^ cells/mL in 96-well plates and bacteria were grown O/N in LB liquid media from a frozen stock. OD_600_ was measured the next day and the appropriate MOI was adjusted directly in cell culture media for infection. A centrifugation step at 188*×*g 5 min allowed to synchronize bacterial entry before incubating for 1h. Cells were then washed once and incubated in cell media with gentamicin (epithelial cells: 20 µg/ml ; macrophages: 10 µg/ml) for extracellular bacteria killing. At 1 h post-gentamicin treatment for invasion assays, or at different time points post-treatment for intracellular survival assays (3 h, 7 h, 23 h), cells were washed once in PBS and lysed in distilled water. Intracellular bacteria were serially diluted, plated on LB agar plates, and incubated at 28°C for 48 h.

### Immunofluorescence microscopy

Infection assays were performed as described in the previous section, using 24-well plates with coverslips. At different times post-gentamicin treatment (4 h and 23 h), cells were fixed with 4% paraformaldehyde for 15 min, washed with 1% bovine serum albumin (BSA) in PBS, permeabilized in PBS supplemented with 0.1% triton (100X)-1% BSA for 4 min and washed in 1% BSA-PBS. Primary antibodies (α-O1b, α-LAMP1) or secondary antibodies with Hoechst (listed in Table 4) all diluted in 1 % BSA were added for 30 min labelling at room temperature in two independent steps with washes in between. After washings, coverslips were fixed on a slide (Fluoromont-G, Invitrogen), and analyzed with an epifluorescent microscope (Zeiss Axiovert 200) using the image acquisition software Metamorph (Molecular Devices). Images were treated with Fiji software.

**Table 4:** Antibodies used for immunofluorescence staining.

| Antibody and dye | Brand and dilution |
| --- | --- |
| Rabbit anti-O1b of *Y. pseudotuberculosis* | CNR stock (1/500<sup>e</sup>) |
| Mouse anti-human LAMP-1, clone H4A3 | Developmental Studies Hybridoma Bank (1/500<sup>e</sup>) |
| Rat anti-mouse LAMP-1, clone 1D4B | Developmental Studies Hybridoma Bank (1/500<sup>e</sup>) |
| Goat anti-rabbit Alexa Fluor 546 | Invitrogen, Thermo Scientific (1/200<sup>e</sup>) |
| Goat anti-mouse Alexa Fluor 488 | Invitrogen, Thermo Scientific (1/200<sup>e</sup>) |
| Goat anti-rat IgG (H+L), Alexa Fluor 488 | Invitrogen, Thermo Scientific A-11006 (1/200<sup>e</sup>) |
| Goat anti-mouse IgG (H+L), Alexa Fluor 568 | Invitrogen, Thermo Scientific A-11004 (1/1000<sup>e</sup>) |
| Hoechst | Invitrogen, Thermo Scientific (1/1000<sup>e</sup>) |

### Transmission electron microscopy

HeLa cells were seeded in a 6 well-plate at a density of 2*×*10^6^ cell per well in quadruplicate. Infection was performed as described in the gentamicin protection assay section (see above). At different times post-gentamicin treatment (1 h, 4 h and 23 h), cells were washed with PBS and detached with trypsin (Gibco). The cell suspension was washed once in PBS and fixed with a 4% PFA, 2.5% glutaraldehyde PBS buffer and kept at 4°C before treatment. Samples were then washed in phosphate-buffered saline (PBS) and post-fixed by incubation with 2% osmium tetroxide (Agar Scientific, Stansted, UK) for 1 h. Samples were then fully dehydrated in a graded series of ethanol solutions and propylene oxide. Impregnation step was performed with a mixture of (1:1) propylene oxide/Epon resin (Sigma) and then left overnight in pure resin. Cells were then embedded in Epon resin (Sigma), which was allowed to polymerize for 48 hours at 60°C. Ultra-thin sections (90 nm) of these blocks were obtained with a Leica EM UC7 ultramicrotome (Wetzlar, Germany). Sections were stained with 5% uranyl acetate (Agar Scientific), 5% lead citrate (Sigma) and observations were made with a transmission electron microscope (JEOL 1011, Tokyo, Japan).

### Mice infection

Female 7-weeks-old BALB/cByJ mice (Charles River France) were acclimatised to the Institut Pasteur animal facility for one week prior to experimentation. For inoculum preparation, bacteria were grown overnight in LB liquid at 28°C and transferred on LB agar plate for 24 h at 28°C. Bacteria on plate were resuspended in PBS and adjusted to OD_600_ = 0.23 to reach a dose of 6*×*10^6^ bacteria per mouse in 20 µL of PBS. Oral infection was performed using the bread feeding method previously described (Derbise et al., 2020). For intravenous (IV) infection, bacterial suspensions were prepared in the same way as described above but serially diluted to reach around 10, 50 or 90 bacteria in 100 µL of PBS. For IV injection, the mouse tail was heated with a paper soaked in warm water prior to injection in the caudal vein with a 29G needle. Inocula were confirmed by plating serial dilutions or directly the IV inoculum on LB agar plates. After infection, animals were monitored for clinical state and weighted daily for 22 days. All experiments were performed in accordance with the Institut Pasteur’s guidelines for laboratory animal welfare.

### Evaluation of the bacterial load in mouse infected organs

For determination of bacterial colonization, organs were collected on euthanized mice at specific days post-infection (5 and 7) and homogenized in PBS in appropriate tubes depending on the organ. cæcum, spleen and a pool of mesenteric lymph nodes were treated in glass beads-containing tubes following two cycles of 30 s with 60 s of “pause” with an homogenizer (Precellys24, Bertin Instruments). Intestines and colons were first opened longitudinally for content collection, then washed thrice in DMEM before being treated for one hour in DMEM supplemented with gentamicin (40 µg/mL) at room temperature. Livers, intestines and colons were disrupted using gentleMACS M tubes (Miltenyi Biotec) with the preloaded program “m_heart_02” on gentleMACS Dissociator (Miltenyi Biotec). Intestinal and colon contents were homogenized using disposable homogenizers (Kimble Chase piston pellet, Fisher Scientific). Serial dilutions of organ homogenates were plated on LB agar with irgazan and incubated at 28°C for 48 h. To avoid contamination from cæcum, intestinal and colon contents plates were grown for a week at 4°C.

### Cytokine measurement

Cytokine concentrations were measured using the mouse magnetic 18-plex Luminex assay (Biotechne, R&Dsystems) on supernatants from infected mBMDM cells. The complete list of cytokines and chemokines is listed in Table 5. Cell supernatants were collected at 8 h post-infection, centrifuged at 5 000*×*g for 5 min at 4°C in SpinX 0,2 µm tubes (Corning, Fisher Scientific) for cell debris and bacterial removal, and stored at −80°C. Samples were processed following the manufacturer’s instructions of the Luminex kit coupled to the DropArray technology (Curiox, Biosystems) which include: the LT washing station MX, DA-bead 96-well, Curiox humid box with Magnet array. Washing steps were performed in 0.1% BSA, 0.05% Tween20 in PBS and the blocking of the DA-bead plate in 10% BSA solution.

**Table 5:** List of cytokines and chemokines in the 18-plex bead immunoassay luminex kit.

| Cytokines and chemokines complete name |
| --- |
| CXCL1/GRO-α/KC/CINC-1 |
| CXCL2/GRO-β/MIP-2/CINC-3 |
| CXCL10/IP-10/CRG-2 |
| CCL2/JE/MCP-1 |
| CCL3/MIP-1α |
| CCL4/MIP-1β |
| CCL5/RANTES |
| G-CSF |
| IFN-γ |
| IL-1α/IL-1F1 |
| IL-1β/IL-1F2 |
| IL-2 |
| IL-4 |
| IL-6 |
| IL-7 |
| IL-10 |
| IL-13 |
| IL-33 |

### Immunohistochemistry and analyses

Cæcum and pools of mesenteric lymph nodes were collected from bread-infected mice or non-infected mice (PBS) at 7 days post-infection, placed in tissue cassettes (Leica Biosystems) and rapidly plunged into formol for 24 h fixation. Samples were then put in a 70% ethanol solution before treatment. Samples were fixed in 10% neutral buffered formalin for 14 hours and embedded in paraffin. Paraffin sections (5-µm thick) were processed for routine histology using hematoxylin and eosin (HE). Stained slides were scanned with ZEISS Axio Scan 7 Digital Slide Scanner and evaluated with Zen software (Zeiss). Histopathological evaluation of HE samples were blindly examined by two pathologists. In each cæcum samples, histopathological criteria summarized in Table 6 were applied. For MLNs, criteria were evaluated as reported in (El-Maraghi and Mair, 1979) with findings as followed: lymphoid hyperplasia, apoptosis, histiocytic hyperplasia, granulomas, necrotizing granulomas and scored with degree (0) absence, (1) mild, (2) moderate, (3) severe. The sum of all criteria represents the total microscopic score. HE scanned slides were subjected to histomorphometry to calculate the total percentage of necrosis area in each sample by using Nikon NIS-Ar software (Nikon Instruments Inc., NY, United States). In addition, serial sections were prepared for immunohistochemistry (IHC) analyses. After a 20 minutes at 100°C pH 6 Citrate buffer antigen retrieval, an IHC was performed by using the Bond RX robot (Leica) with rabbit anti-F4/80 antibody (D2S9R, Cell Signaling) or rat anti-Ly6B2 (MCA77, biotinylated G, BioRad), goat anti-rabbit Ig secondary antibody (E0432, Dako, Agilent) or rat IgG-HRP secondary antibody (VC005-025, R&D Systems) and Leica Streptavindine-HRP/DAB detection Kit. IHC stained slides (F4/80 and Ly2B6) were scanned (Hamamatsu S360, Hamamatsu Photonics, Shizuoka, Japan) and subjected to digital image analysis to calculate the percentage of positivity labeling with respect to the whole area of the organ by using Nikon NIS-Ar software (Nikon Instruments Inc., NY, United States).

**Table 6:** Histological criteria applied in cæcum samples.

| **Finding** | **Definition** | **Degree** | **Extension** |
| --- | --- | --- | --- |
| *Inflammatory cell infiltrate* |  |  |  |
| Leukocyte cell infiltration | Leukocyte infiltration in the mucosa or lamina propria, sometimes forming leukocyte aggregates. | Absence (0), Mild (1), Moderate (2), Severe (3) | Focal (1), Multifocal (2), Diffuse (3) |
| Granulomas | Epithelioid macrophages infiltration in the mucosa or lamina propria | Absence (0), Mild (1), Moderate (2), Severe (3) | Focal (1), Multifocal (2), Diffuse (3) |
| *Mucosal architecture* |  |  |  |
| Necrosis | Necrosis with presence of free bacteria, prominent acute inflammation (mainly, neutrophils) haemorrhage and loss of mucosal architecture | Absence (0), Mild (1), Moderate (2), Severe (3) | Focal (1), Multifocal (2), Diffuse (3) |
| Submucosal oedema | Connective tissue areas of lower optical density below the mucosal layer | Absence (0), No inflam. (1), inflam. (2) | Focal (1), Multifocal (2), Diffuse (3) |
| Mucosal destruction | Loss of mucosal architecture with necrosis of epithelium | Absence (0), Presence (1) | Focal (1), Multifocal (2) |
| *Epithelial changes* |  |  |  |
| Erosion of the epithelium | Loss of surface epithelium | Absence (0), Presence (1) | Focal (1), Multifocal (2) |
| Cryptitis | Neutrophils between crypt epithelial cells | Absence (0), Presence (1) | Focal (1), Multifocal (2) |
| Crypt abscesses | Neutrophils and cell debris in crypt lumen | Absence (0), Presence (1) | Focal (1), Multifocal (2) |
| Goblet cell hyperplasia | Increase in the number of goblet cells | Absence (0), Presence (1) | Focal (1), Multifocal (2) |

### Statistical analyses

Statistical analyses were performed using GraphPad Prism (GraphPad Prism 10.2.2 Software). Statistical significance is expressed as follow: *P<0.05, **P<0.01, ***P<0.001, ****P<0.0001. Survival data were plotted using the Kaplan-Meier estimator and log- rank (Mantel-Cox) tests were used to assess differences in survival between groups. Cytokine concentrations were transformed in log(Y+1) before computed for a two-way ANOVA (or mixed model) using Prism.

## Acknowledgements

We are grateful to all members of the *Yersinia* Research Unit and the *Yersinia* National Reference Laboratory for helpful comments, insightful discussions and particularly Jose Pablo Marín Obando for his contribution to parallel works linked to this project. We thank the Histopathology core Facility at Institut Pasteur for sample processing for histology and immunohistochemistry, and more specifically David Hing for the entire technical support and David Hardy for project management support. We thank the members of the Cytometry and Biomarkers UTechS Platform (CB UTechS) at Institut Pasteur for technological support, and specifically Esma Karkeni for training and advice. We also acknowledge the help of Institut Pasteur Central animal facility team for their support with mouse hosting and care. This work received financial support from Institut Pasteur, Université Paris Cité and LabEX Integrative Biology of Emerging Infectious Diseases (ANR-10_LBX-62-IBEID). ML received a Ph.D. fellowship from MENESR (Ministère de l’Education Nationale, de l’Enseignement Supérieur et de la Recherche). We declare no conflict of interest.

## Supplementary figures and tables

**Figure S1:**
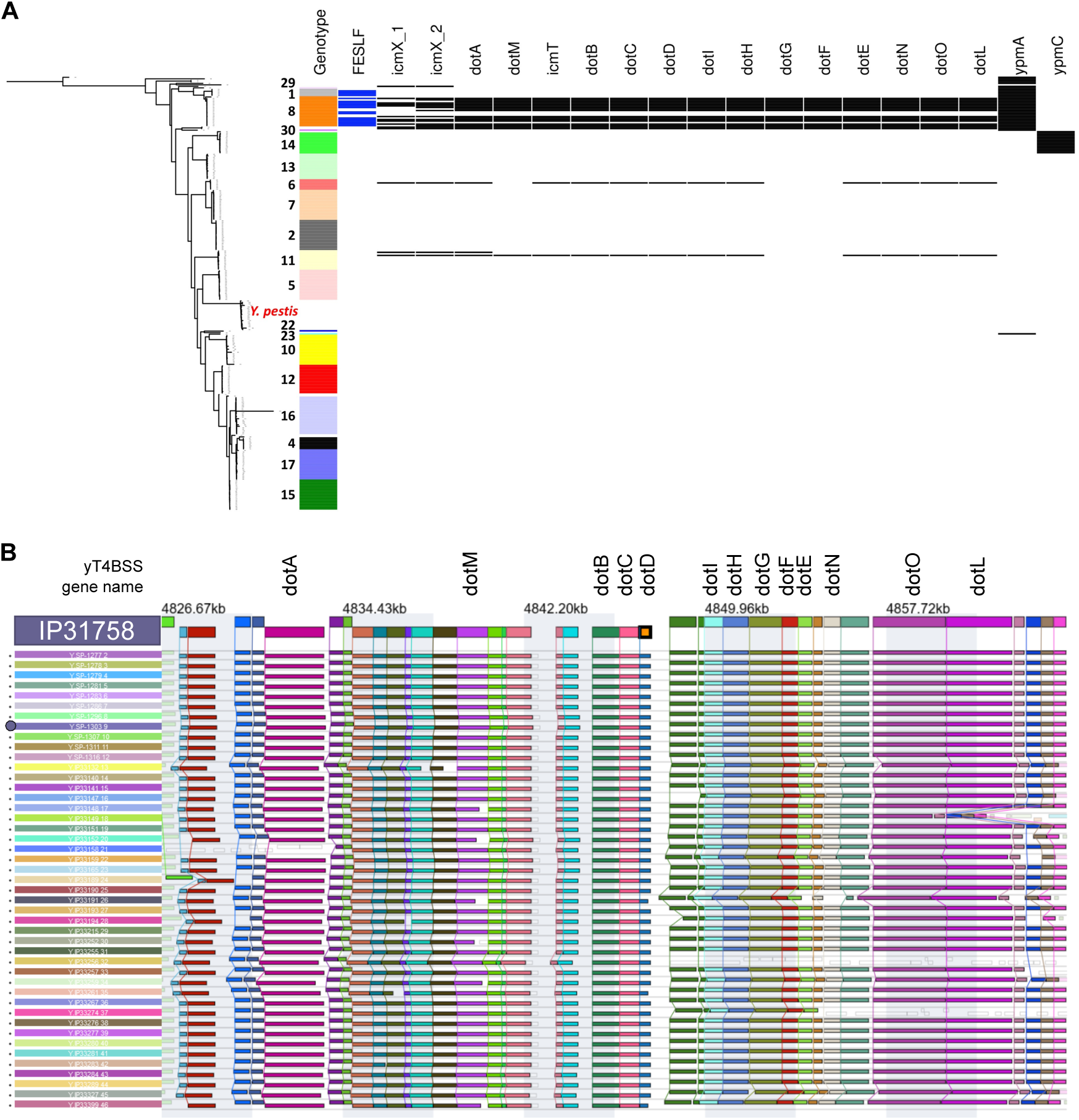
*y*T4BSS genes are clustered in *Y. pseudotuberculosis* lineage 8 and display a conserved synteny. **(A)** Presence of *y*T4BSS and *ypm* genes along the phylogenetic tree of *Y. pseudotuberculosis* performed on 287 genomes (including 20 *Y. pestis* isolates). Genes coloured in black correspond to the presence and in white absence of genes using a threshold of 80 % identity and 80 % coverage. The strain SP-1303 was taken as reference strain. Strains with clinical data associated with FESLF are coloured blue. **(B)** Synteny analysis of the *y*T4BSS operon between 46 *Y. pseudotuberculosis* genomes from lineage 8. The strain IP31758 is used as a reference for this analysis. The purple star highlights the SP-1303 strain used as a reference in this study. Genes are aligned on *dotD*. The synteny analysis was performed with SynTView (Lechat et al., 2013).

**Figure S2:**
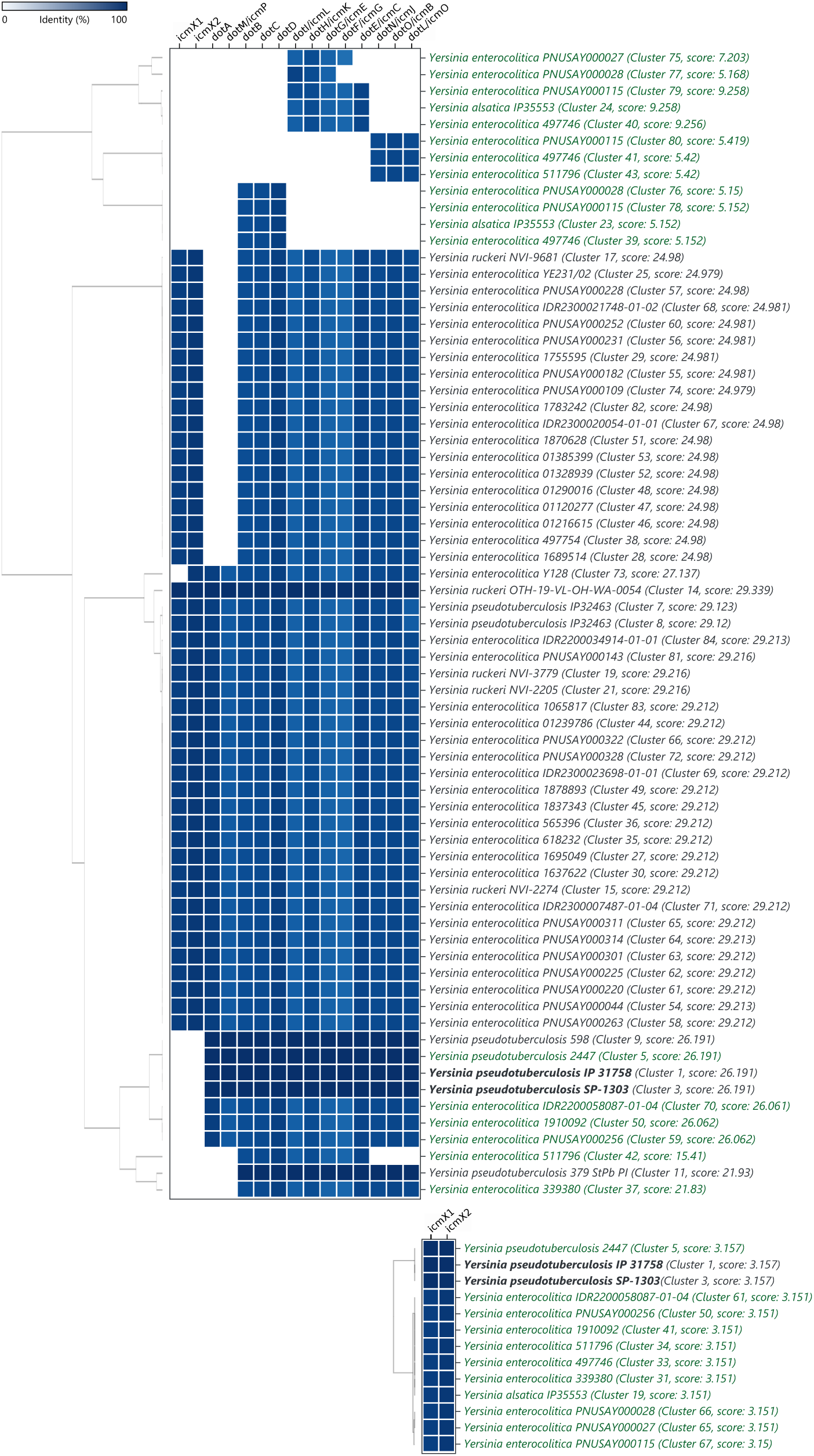
Homology of the *y*T4BSS gene cluster of *Y. pseudotuberculosis* SP-1303 with *Yersinia.* spp. Analysis was computed with cblaster against NCBI genome database, with default parameters: minima of 50% query coverage and 30% identity (Gilchrist and Chooi, 2021).

**Figure S3:**
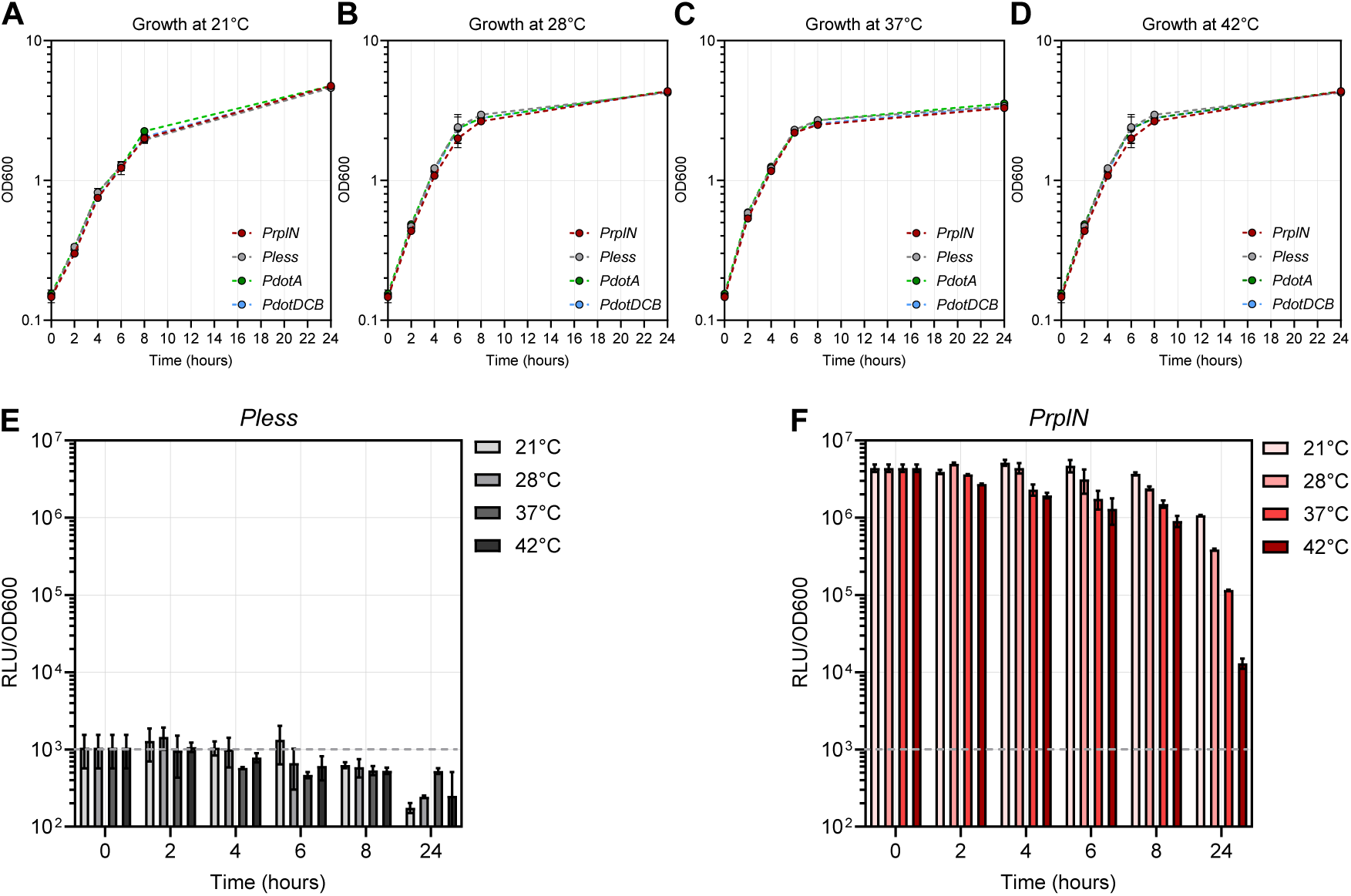
Growth condition of bioluminescent reporters and bioluminescent emission of a constitutive bioluminescent reporter *PrplN-lux* and a control strain without promoter *Pless-lux* in LB media under different temperatures. **(A-D)** Growth curve of the different *Y. pseudotuberculosis* strains harboring bioluminescent reporters in LB media at 4 temperatures tested (21°C, 28°C, 37°C, 42°C). Data represent the mean and SD of two independent experiments. **(E-F)** Measurement of photon emission produced during growth in LB media of control bacterial strains harboring bioluminescent reporters at different temperatures (21°C, 28°C, 37°C, 42°C) over time. *Pless-lux* does not harbour any promoter and is used as a negative control for signal background while *PrplN-lux* is used as a constitutive promoter. Bacteria were grown O/N at 28°C prior the experiment. Relative light unit (RLU) measured was normalized by OD_600nm_. Data are representative of two independent experiments with means and SD.

**Figure S4:**
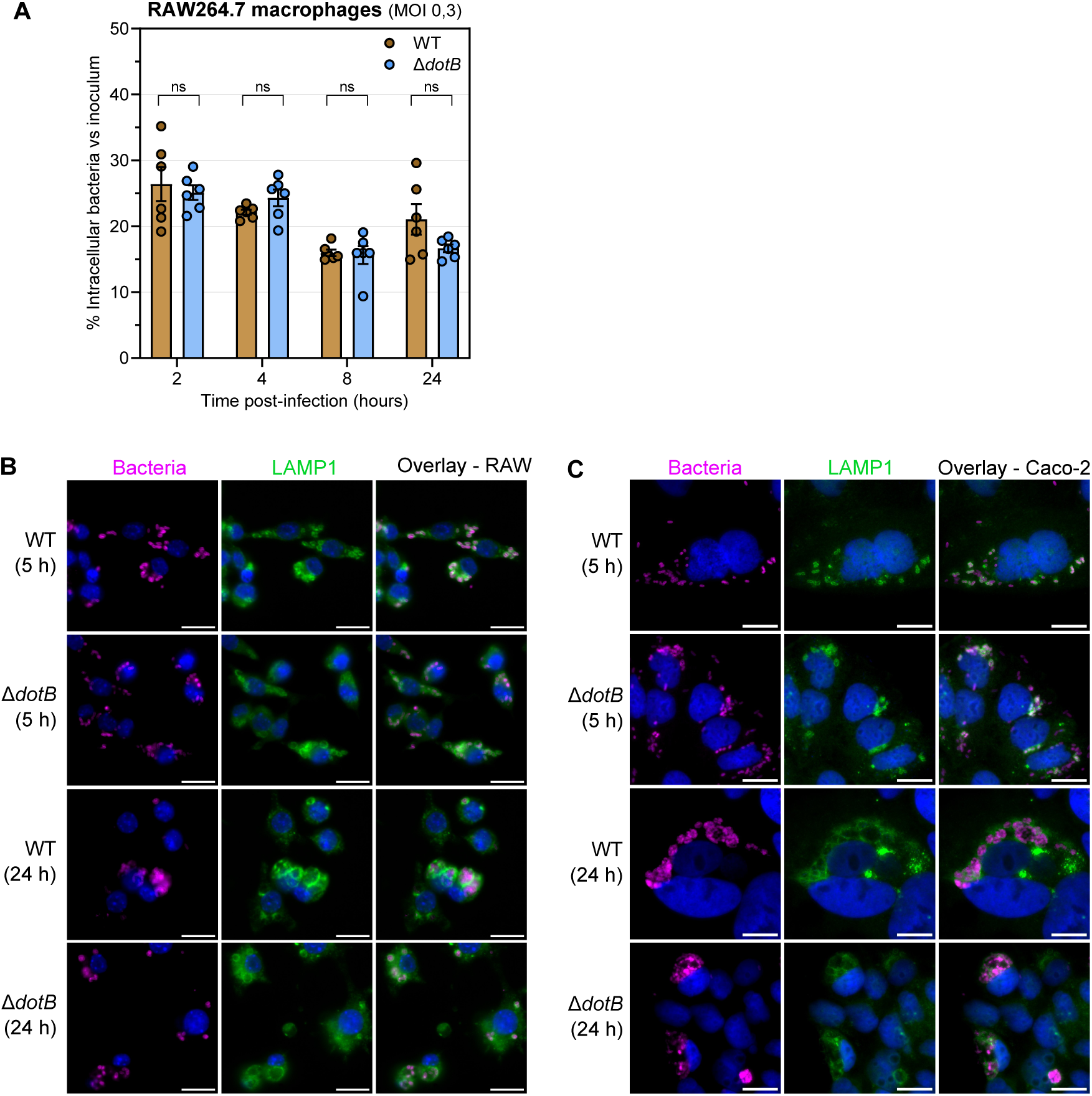
The *y*T4BSS does not contribute to the intracellular life cycle of *Y. pseudotuberculosis* FESLF strain within Caco-2 epithelial cells and Raw264.7 macrophages. **(A)** Intracellular survival between the WT and Δ*dotB* mutant strains within RAW264.7 (MOI 0,3) over time (2 hpi, 4 hpi, 8 hpi, 24 hpi). Data are representative of single experiments with means and SD. Multiple unpaired t tests were performed. **(B)** Representative images of RAW264.7 (MOI10) **(C)** Caco-2 cells (MOI 25), with the WT or Δ*dotB* mutant strains at 5 hpi or 24 hpi by fluorescent using antibodies against surface O-antigen (O1b) of *Y. pseudotuberculosis* (Cy5, magenta), antibodies against LAMP1 (FITC, green) and Hoechst (blue). Scale bars are 10 µm. Overlay images were generated using ImageJ software.

**Figure S5:**
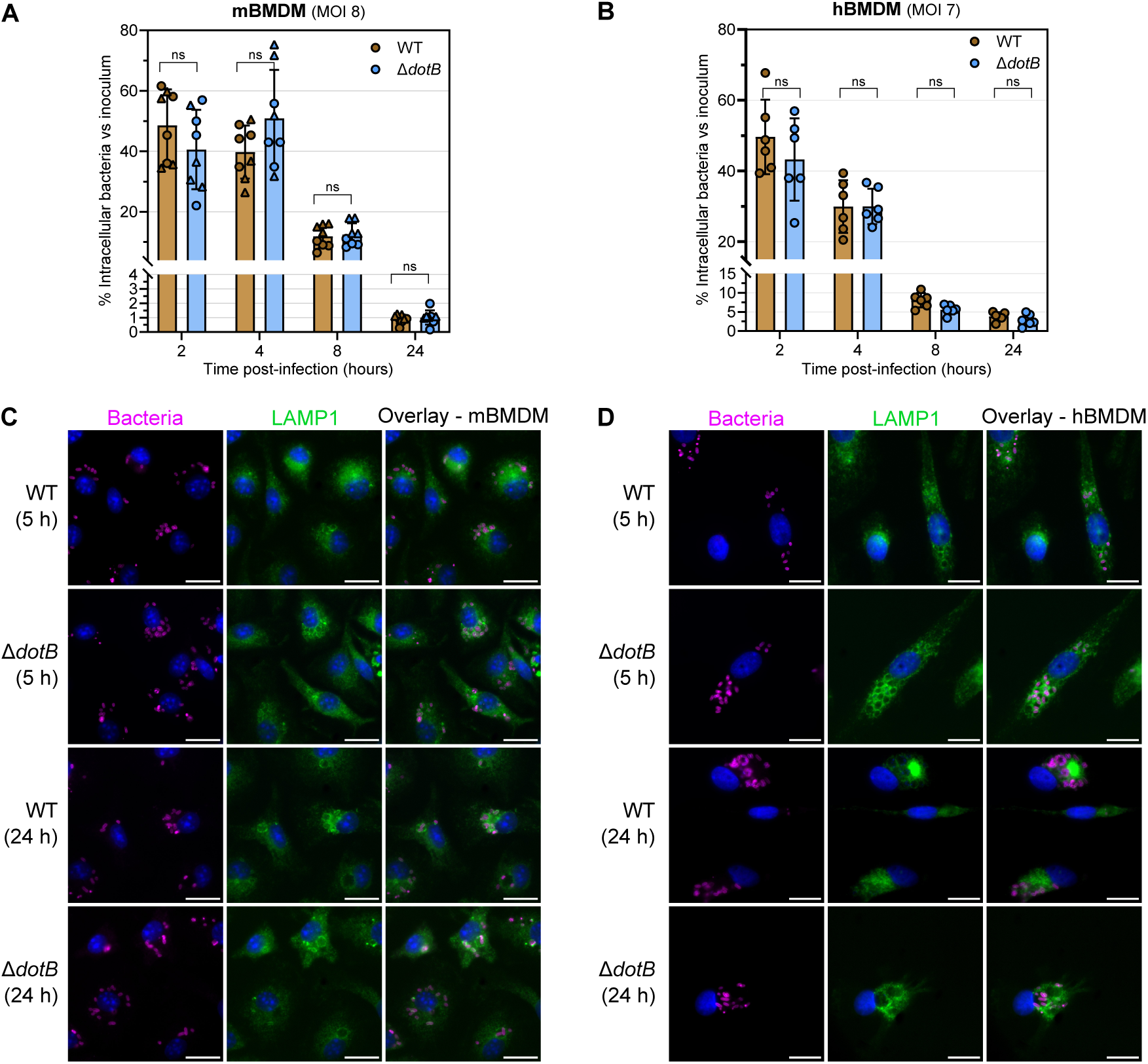
The *y*T4BSS does not contribute to the intracellular life cycle of *Y. pseudotuberculosis* FESLF strain within mBMDM and human blood monocyte-derived macrophages. **(A-B)** Intracellular survival between the WT and Δ*dotB* mutant strains within mBMDM (MOI 8) and human blood monocyte-derived macrophages (MOI 7) over time (2 hpi, 4 hpi, 8 hpi, 24 hpi). Data are representative of two independent experiments (triangles and circles differentiate the experiments) with means and SD. Multiple unpaired t tests were performed. **(C-D)** Representative images of mBMDM, human blood monocyte-derived macrophages infected (MOI 10) with the WT or Δ*dotB* mutant strains at 5 hpi or 24 hpi by fluorescent using antibodies against surface O-antigen (O1b) of *Y. pseudotuberculosis* (Cy5, magenta), antibodies against LAMP1 (FITC, green) and Hoechst (blue). Scale bars are 10 µm. Overlay images were generated using ImageJ software.

**Figure S6:**
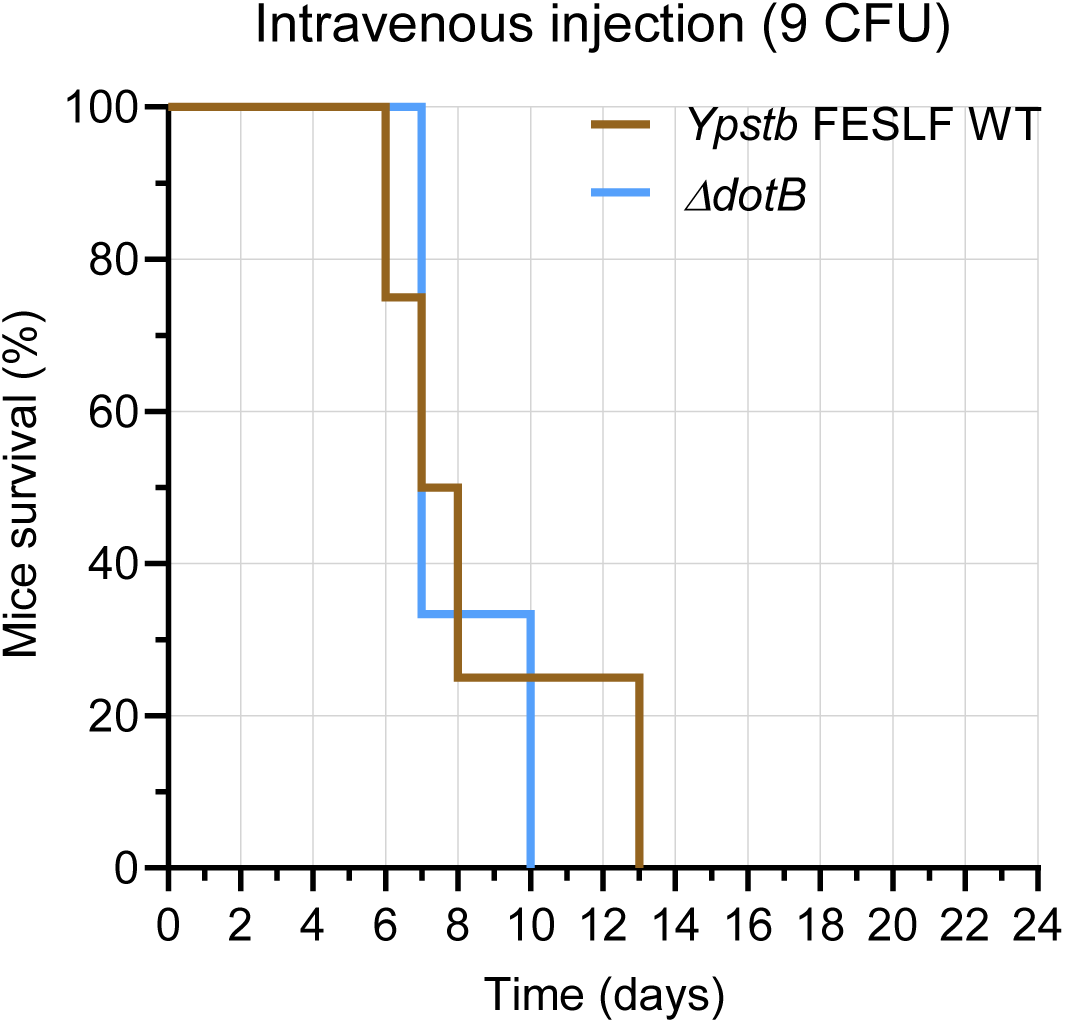
The *y*T4BSS does not participate in virulence of *Y. pseudotuberculosis* upon intravenous infection of mice. Survival monitoring of mice after intravenous injection of 9 CFU/mouse. Representation of a single experiment (n=4/condition).

**Figure S7:**
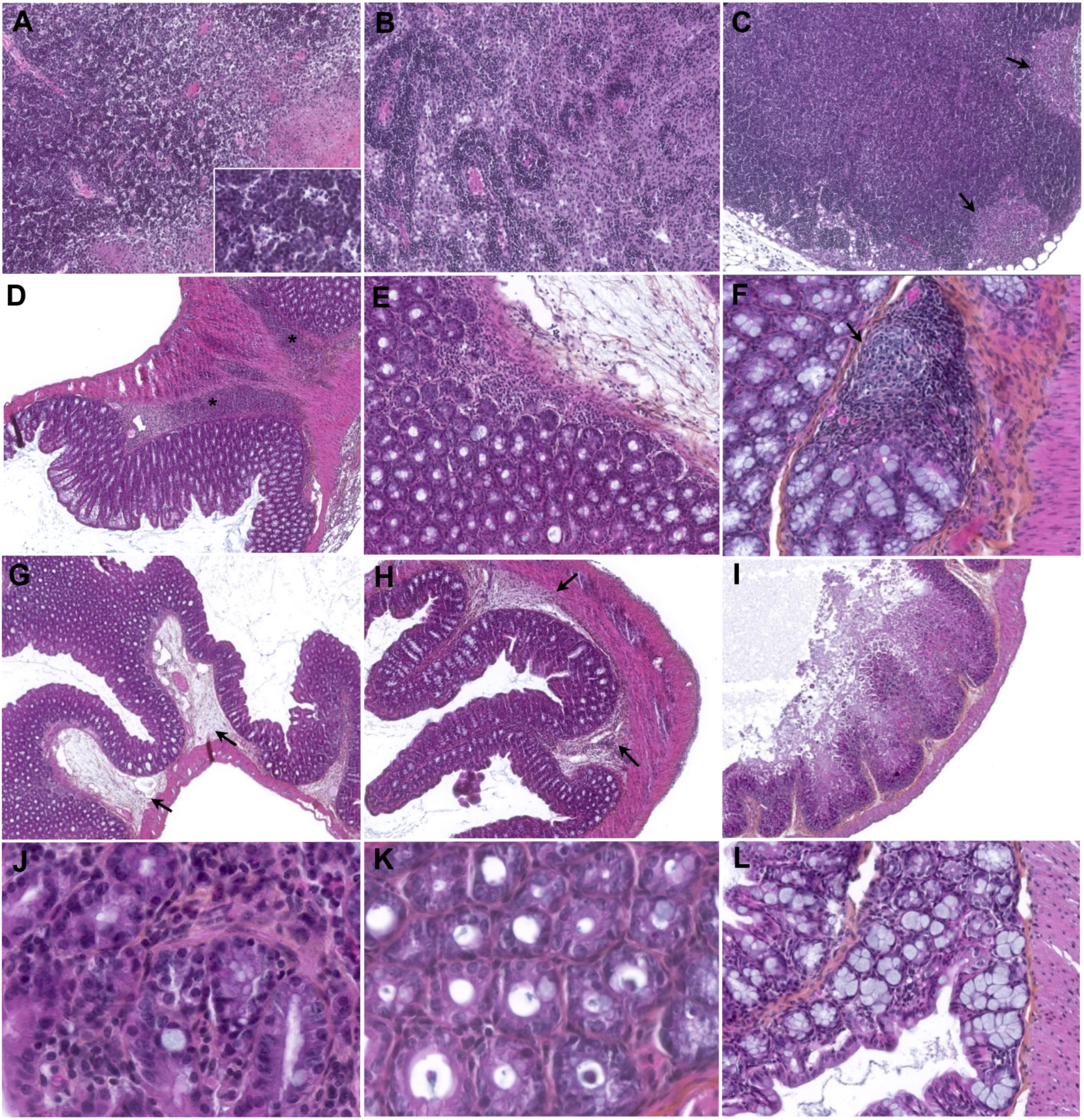
Other histopathological finding in MLNs and cæcum observed in infected mice (H&E). **(A-C)** Representative pictures of histopathological findings found in MNL from infected mice. **(A)** Moderate level of apoptosis in the MLNs from a Δ*dotB*-infected mouse. Inset show tingible body macrophages. **(B)** Histiocytic hyperplasia in the MLNs from a WT-infected mouse. **(C)** Granulomas (arrows) in the MLNs from Δ*dotB*-infected mouse. **(D-L)** Representative pictures of histopathological findings found in cæcum from infected mice. **(D)** Diffuse inflammatory infiltration in the lamina propia from a Δ*dotB*-infected mouse. **(E)** Higher magnification of **(D)**. **(F)** Granuloma in the lamina propria from a WT-infected mouse. **(G)** Oedema in lamina propria (arrows) from a Δ*dotB*-infected mouse. **(H)** Oedema in lamina propria (arrows) from a with severe inflammation from a Δ*dotB*-infected mouse. **(I)** Severe mucosal destruction from a Δ*dotB*-infected mouse. **(J)** Cryptitis from a Δ*dotB*-infected mouse. **(K)** Crypt abscesses from a Δ*dotB*-infected mouse. **(L)** Goblet cell hyperplasia from a Δ*dotB*-infected mouse.

**Figure S8:**
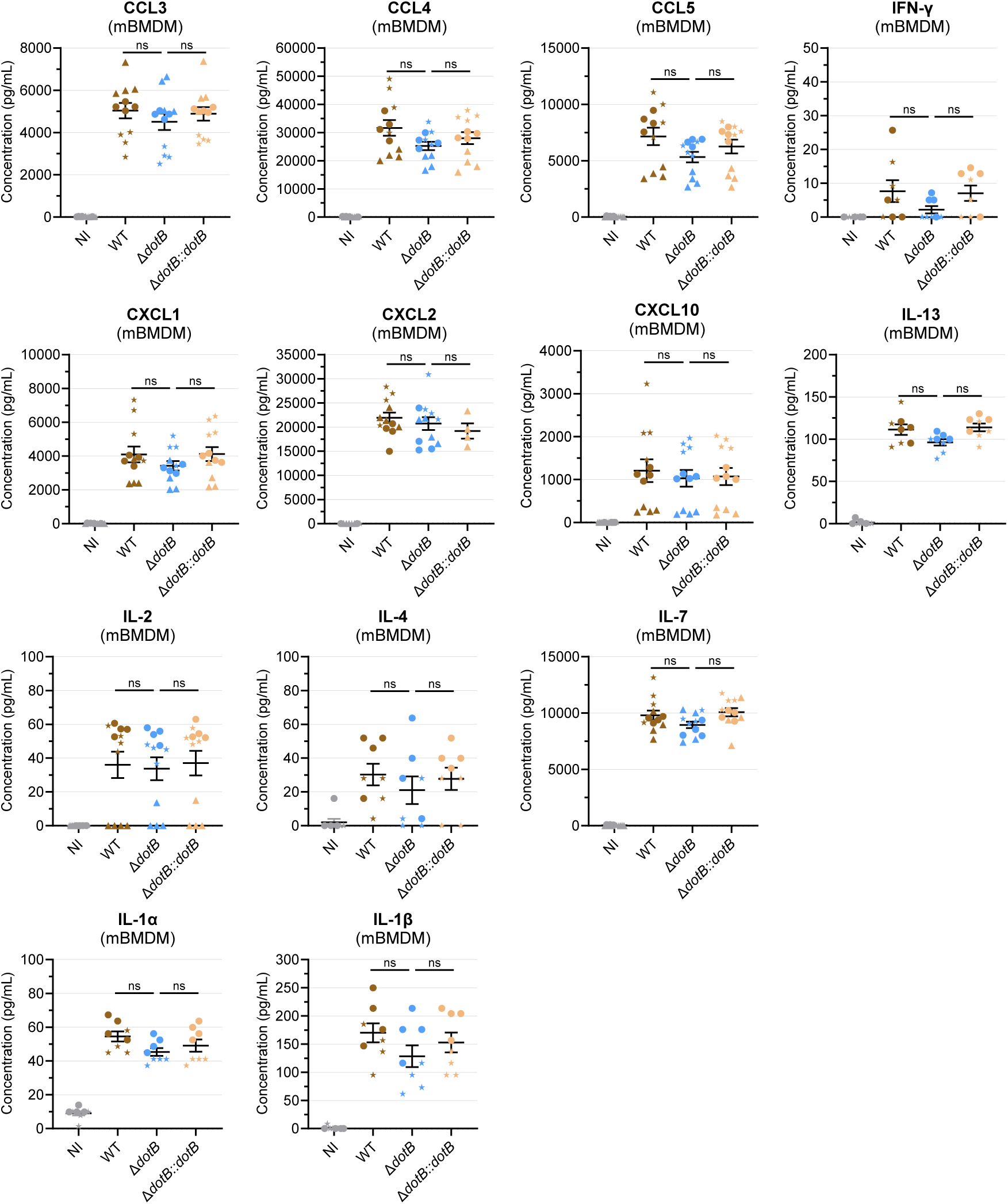
Panel of explored cytokines from supernatants of infected mBMDM not modulated by the *y*T4BSS. Cytokine profile analysis on supernatants from infected mBMDMs at 8 hpi, with WT, Δ*dotB* mutant or Δ*dotB::dotB* complemented strains (MOI 10) using Luminex technology. Data are representative of 3 independent experiments with 4 technical replicates and presented as mean and SEM. Mixed model with Tukey’s multiple comparisons correction were performed on logY+1 transformed data.

